# Type II alveolar cells with constitutive expression of MHCII and limited antigen presentation capacity contribute to improved respiratory viral disease outcomes

**DOI:** 10.1101/2021.03.18.435984

**Authors:** Sushila A. Toulmin, Chaitali Bhadiadra, Andrew J. Paris, Jeffrey H. Lin, Jeremy Katzen, Maria C. Basil, Edward E. Morrisey, G. Scott Worthen, Laurence C. Eisenlohr

**Affiliations:** Division of Protective Immunity, Department of Pathology and Laboratory Medicine, Children’s Hospital of Philadelphia, Philadelphia, PA, USA; Perelman School of Medicine, University of Pennsylvania, Philadelphia, PA, USA; Division of Pulmonary, Allergy and Critical Care Medicine, Department of Medicine, Perelman School of Medicine, University of Pennsylvania, Philadelphia, PA, USA; Department of Medicine, Penn-CHOP Lung Biology Institute, University of Pennsylvania, Philadelphia, PA, USA; Penn Cardiovascular Institute, Perelman School of Medicine, University of Pennsylvania, Philadelphia, PA, USA; Penn Institute for Regenerative Medicine, Perelman School of Medicine, Philadelphia, PA, USA; Department of Cell and Developmental Biology, Perelman School of Medicine, University of Pennsylvania, Philadelphia, PA, USA; Department of Pediatrics, University of Pennsylvania Perelman School of Medicine, Philadelphia, PA, USA; Division of Neonatology, Children’s Hospital of Philadelphia, Philadelphia, PA, USA; Department of Pathology and Laboratory Medicine, Perelman School of Medicine, University of Pennsylvania, Philadelphia, PA, USA

## Abstract

Type II alveolar cells (AT2s) are critical for basic respiratory homeostasis and tissue repair after lung injury. Prior studies indicate that AT2s also express major histocompatibility complex II (MHCII) molecules, but how MHCII expression by AT2s is regulated and how it contributes to host defense remain unclear. Here we show that AT2s express high levels of MHCII independent of conventional inflammatory stimuli, and that selective loss of MHCII from AT2s in mice results in the worsening of respiratory virus disease following influenza and Sendai virus infections. We also find that AT2s exhibit MHCII presentation capacity that is substantially limited in comparison to professional antigen presenting cells. The combination of constitutive MHCII expression and restrained presentation may position AT2s to contribute to lung adaptive immune responses in a measured fashion, without over-amplifying damaging inflammation.

## Main

CD4^+^ T cell responses are initiated by T cell receptor (TCR) recognition of cognate major histocompatibility complex II (MHCII)/peptide complexes on the surface of antigen presenting cells (APCs). By convention, constitutive MHCII expression is restricted to a few specialized types of cells, including the “professional APCs” B cells, dendritic cells (DCs), and macrophages, as well as thymic epithelial cells (TECs)^1^. In contrast, all other cell types are thought to produce MHCII only in the setting of inflammation, specifically in response to the cytokine interferon-γ (IFNγ)^2^. However, more recently, an increasing number of other cell types have been shown to express MHCII at homeostasis and to contribute to adaptive immune responses in a variety of settings^3, 4^. These include certain intestinal ILC3 subsets^5, 6^, which express MHCII independent of IFNγ, as well as several nonhematopoietic cell types, such as lymph node stromal cells^7–10^, vascular endothelial cells^11–13^, and intestinal epithelial cells^14–16^, whose MHCII production is driven by the basal IFNγ present at steady state.

Type II alveolar cells (AT2s) are an additional nonhematopoietic cell type that have been reported to express MHCII at homeostasis in both rodents^17–22^ and humans^23–27^. AT2s are epithelial cells present in the lung parenchyma, and their main physiologic functions are to produce surfactant and to serve as stem-like cells in the distal lung^28–31^. AT2s have also been described to perform several innate immunologic functions^32–34^, and their constitutive MHCII expression suggests that they may participate in lung adaptive immune responses as well. Previous studies of AT2 MHCII function have varied widely, with some concluding that AT2s serve to activate CD4^+^ T cells^19, 21^ and others demonstrating the opposite^20, 26^. As a result, both the MHCII presentation capability of AT2s and the contribution of AT2 MHCII to lung host responses remain unclear. Here we report on the mechanisms regulating homeostatic MHCII expression in AT2s, their antigen presentation capacity, and the impact of AT2 MHCII *in vivo*.

## Results

### AT2s express MHCII protein at homeostasis to uniformly high levels

To verify prior reports of homeostatic AT2 MHCII expression, we assessed MHCII on both mouse and human AT2s by flow cytometry. We found that naïve wild-type C57Bl/6 (B6) AT2s, identified as CD45^-^CD31^-^EpCAM^int^ (Fig S1)^22^, uniformly express MHCII (Fig 1a). Similarly, healthy human HT2-280^+^ AT2s^35^ express the human MHCII protein HLA-DR (Fig S2a,b).

**Figure 1:**
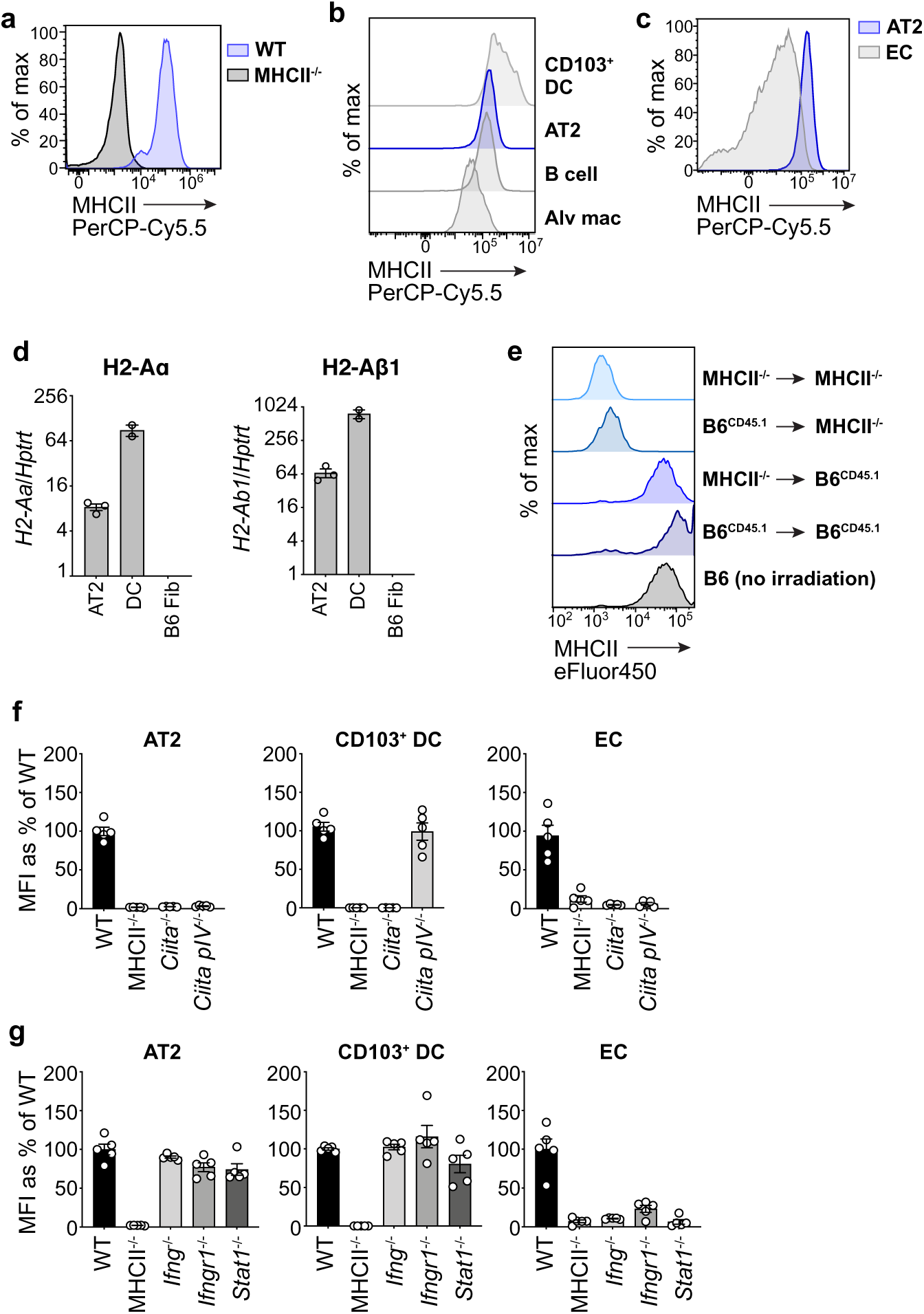
Type II alveolar cells (AT2s) constitutively express MHCII independent of conventional inflammatory signals. **a-c**, MHCII protein expression by AT2s from naïve C57Bl/6 (B6) wild-type (WT) and MHCII^-/-^ mice (**a**) and AT2s, CD103^+^ dendritic cells (CD103^+^ DC), B cells, alveolar macrophages (Alv mac), and endothelial cells (EC), from naïve B6 WT lungs (**b,c**) measured by *ex vivo* flow cytometry; histograms represent n>10 mice per strain total from >5 independent experiments. **d,** I-A^b^ α (H2-Aα) and β (H2-Aβ1) chain transcript abundance relative to housekeeping gene Hprt1, in AT2s and lung dendritic cells (DC) sorted from naïve B6 WT mice, and a B6 fibroblast cell line (B6 Fib), measured by quantitative RT-PCR; n=3 mice for AT2s, n=2 mice for DCs, with 3 technical replicates for each measurement. **e,** MHCII protein expression by AT2s in MHCII^-/-^ and CD45.1 B6 (B6^CD45.1^) bone marrow chimeric mice, measured 8 weeks post-transplantation with transfers as indicated; histograms represent n=2-4 mice per group in 1 experiment. **f,g,** MHCII protein expression by AT2s, CD103^+^ DCs, ECs from naïve B6 WT, MHCII^-/-^ (**f,g**) *Ciita*^-/-^, *Ciita pIV*^-/-^ (**f**), *Ifng*^-/-^, *Ifngr1*^-/-^, and *Stat1*^-/-^ (**g**) lungs, measured by flow cytometry. For each strain, MHCII expression is displayed as a percentage reflecting the average median fluorescence intensity (MFI) of MHCII on each cell type relative to the MHCII MFI on the same cell type in WT mice. Symbols indicate n=5 mice per strain, pooled from 2-3 independent experiments. 3 of 5 *Ciita pIV*^-/-^ mice used were 8 months old, 1 of 5 *Ifngr1*^-/-^ mice used were 5 months old. **d,f,g,** Data are shown as mean plus standard error of the mean (SEM).

We next compared MHCII levels on AT2s to other well-studied MHCII-expressing lung cells in naïve mice. AT2s express MHCII at a magnitude similar to that of lung professional APCs, dendritic cells (DCs) and B cells, and higher than alveolar macrophages (Fig 1b). Besides AT2s, vascular endothelial cells (ECs) are another nonhematopoietic cell type in the lung that express MHCII at homeostasis^36^. Compared to ECs, AT2 MHCII expression is higher and more homogeneous (Fig 1c).

MHCII protein may be synthesized intracellularly or extrinsically acquired^37^ from conventional APCs. We examined the source of AT2 MHCII by RNA-sequencing analysis of naïve mouse AT2s [data originally generated by Ma *et al*^38^], qPCR, and by creating CD45.1 MHCII^+/+^ (B6^CD^^45.1^) and CD45.2 MHCII^-/-^ B6 (MHCII^-/-^) bone marrow chimeric mice. AT2s express transcript for both the α and β chains of the MHCII allele in B6 mice, I-A^b^ (Fig 1d, S5a), and AT2s retained MHCII protein expression to high levels in MHCII^-/-^ → B6^CD45.1^ mice (Fig 1e). Thus, AT2s synthesize their own MHCII and do not depend on acquisition of MHCII protein from hematopoietic APCs.

In sum, AT2s represent a unique population of nonhematopoietic lung cells that produce MHCII protein at homeostasis uniformly to high levels, similar to that of professional APCs.

### AT2 MHCII expression is regulated in an unconventional, IFNγ-independent manner

The class II transactivator (CIITA) is the master transcription factor that drives MHCII expression^2^. To determine whether AT2 MHCII expression is dependent on CIITA, we assessed MHCII in the lungs of *Ciita*^-/-^ mice. Lack of CIITA results in the complete loss of MHCII expression on AT2s, as well as lung DCs and ECs (Fig 1f). Transcription of CIITA begins at three different promoter sites (pI, pIII, and pIV) depending on cell and context specific stimuli. Recruitment of pI occurs in DCs and macrophages, pIII in B cells and plasmacytoid DCs, and pIV in ILC3s and nonhematopoietic cells^2, 6^. To assess whether AT2 MHCII is dependent on pIV, we evaluated mice lacking just the pIV region within the CIITA locus^39^. AT2s from *Ciita pIV*^-/-^ mice fail to express MHCII, as do ECs, while DCs retain MHCII expression as expected (Fig 1f).

The recruitment of pIV in all nonhematopoietic cells (except for thymic epithelial cells (TECs)) is thought to require induction by the cytokine IFNγ^2^. We evaluated MHCII in mice deficient in IFNγ, IFNγR, and the associated downstream pIV-binding transcription factor, STAT1. AT2s retain MHCII expression in *Ifng*^-/-^, *Ifngr1*^-/-^, *and Stat1*^-/-^ mice in a manner similar to DCs and in contrast to ECs, whose MHCII expression is abrogated in the absence of IFNγ signaling (Fig 1g). AT2s, like DCs, exhibit a small reduction in MHCII expression in *Stat1*^-/-^ mice, indicating that IFNγ signaling may upregulate MHCII expression in AT2s, but it is not required for steady state expression.

Besides IFNγ, no other cytokine has been shown to be sufficient to induce MHCII expression in nonhematopoietic cells. However, given the IFNγ-independent expression of MHCII by AT2s, we also examined the requirement of a broad array of conventional innate and adaptive immune mediators, by assessing *Ifnar1^-/-^*, *Ifnar1^-/-^ x Ifngr1^-/-^*, *Myd88^-/-^*, *Stat6^-/-^*, and germ-free mice. AT2s express MHCII at wild-type levels in all cases (Fig S3), indicating that they also do not require a wide spectrum of other, non-IFNγ inflammatory signals for MHCII expression, including type I IFNs, TLR ligands, IL-1, IL-18, IL-33, Th2 cytokines, and the microbiota.

Altogether, these data demonstrate that AT2 homeostatic MHCII expression is CIITA pIV-dependent, but IFNγ-independent. This same transcriptional configuration has been reported for only two other cell types: TECs^40^, which are present in a primary immune organ, and ILC3s^6^, which are immune cells. To our knowledge it has never been described for a cell outside of the immune system.

### AT2s possess classical APC characteristics and MHCII antigen processing and presentation machinery

We next examined whether AT2s possess mediators of classical MHCII presentation. By convention, this pathway begins with extracellular antigen acquisition followed by proteolysis in the endocytic compartment^41–43^. To assess these steps, we evaluated the capacity of AT2s to catabolize DQ-Ova, a self-quenched conjugate of ovalbumin protein that fluoresces after receptor-mediated endocytosis and proteolytic digestion. AT2s exhibited a temperature-dependent increase in fluorescence when incubated with DQ-Ova, less so than DCs but similarly to B cells, indicating that they are capable of active antigen uptake and degradation (Fig 2a). Antigen degradation is mediated by several key enzymes, including asparagine endopeptidase (AEP), gamma-interferon-inducible lysosomal thiol reductase (GILT), and lysosomal/endosomal cathepsin proteases^44^. AT2s have been previously demonstrated to express cathepsins, in particular Cathepsin H (CtsH), which is critical for surfactant protein processing^45, 46^. By RNA-sequencing analysis, we found that naïve mouse AT2s express transcripts for AEP, GILT, and a wide variety of cathepsin enzymes (Fig S5a). Furthermore, we directly assessed the activity of two cathepsins in AT2s that play major roles in antigen processing: Cathepsin D (CtsD) and Cathepsin L (CtsL). We found that AT2s exhibited readily detectable activity of both CtsD and CtsL, which was abrogated by specific inhibitors of each enzyme (Fig S5b,c).

**Figure 2:**
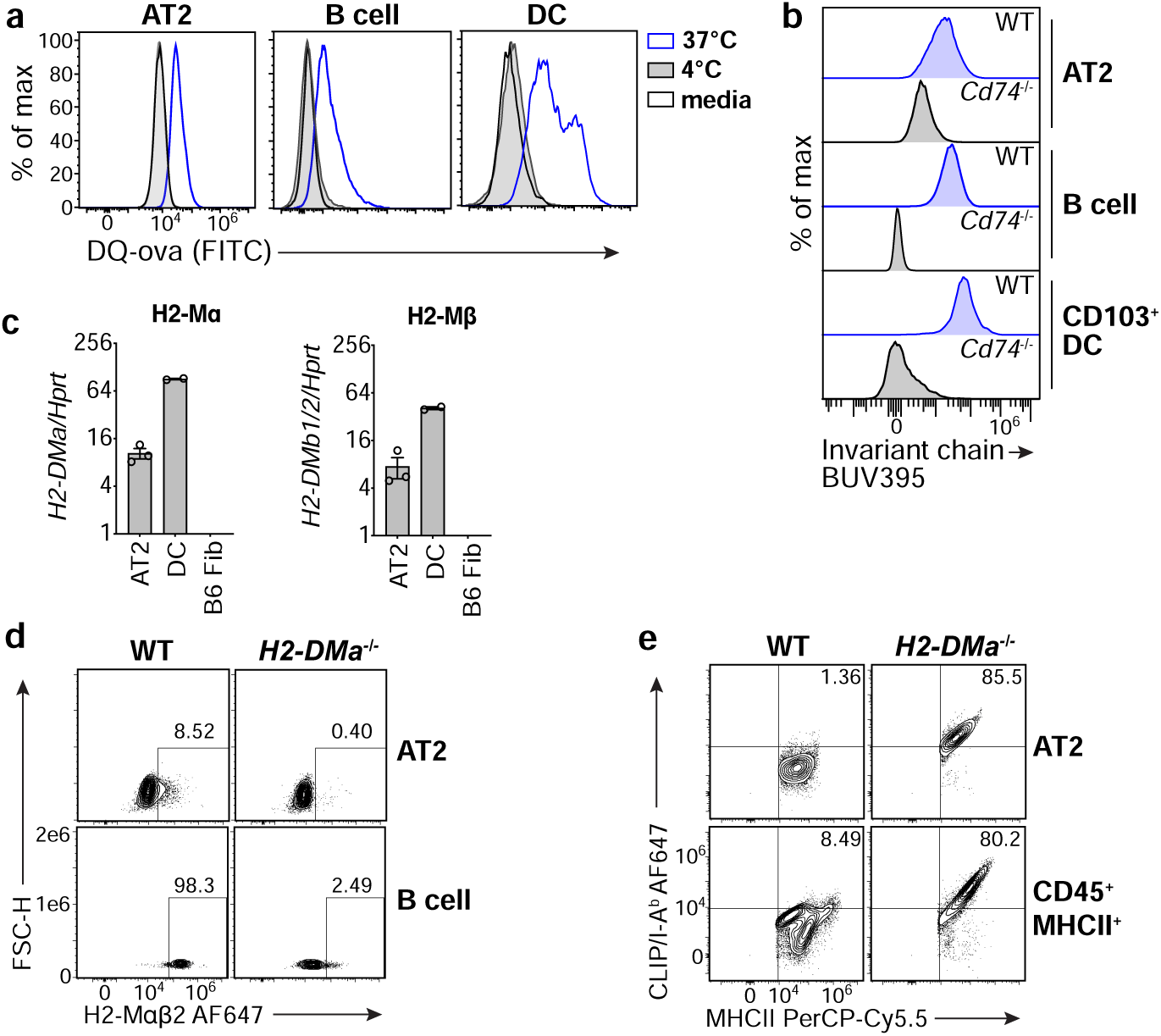
AT2s express canonical MHCII antigen processing and presentation machinery. **a,** Fluorescence of AT2s, B cells, and DCs sorted from naïve B6 WT lungs then incubated with DQ-ovalbumin (DQ-ova) or media at the temperatures indicated, measured by flow cytometry; histograms represent n=2 mice total from 2 independent experiments. **b,** Intracellular invariant chain protein expression by AT2s, lung CD103^+^ DCs, and splenic B cells from B6 WT or invariant chain deficient (*Cd74*^-/-^) mice measured *ex vivo* by flow cytometry; histograms represent n=3-9 mice per strain from 3 independent experiments. **c,** H2-Mα and H2-Mβ chain transcript abundance relative to housekeeping gene Hprt1, in AT2s and DCs sorted from naïve B6 WT lungs, and a B6 fibroblast cell line (B6 Fib), measured by quantitative RT-PCR; n=3 mice for AT2s, n=2 mice for DCs, with 3 technical replicates for each measurement. **d,** Intracellular H2-Mαβ2 protein expression by AT2s and lung B cells from B6 WT or H2-Mα deficient (*H2-DMa*^-/-^) mice measured *ex vivo* by flow cytometry with frequency of H2-Mαβ2^+^ cells shown; plots represent n>10 mice per strain total from >5 independent experiments. Gates were drawn separately for AT2s and B cells based on their H2-Mα deficient cell counterparts. **e,** Surface MHCII and CLIP/I-A^b^ H2-Mαβ2 protein expression by AT2s and bulk CD45^+^MHCII^+^ cells from B6 WT or H2-Mα deficient mice measured *ex vivo* by flow cytometry with frequency of MHCII^+^CLIP/I-A^b+^ cells shown; plots represent n>6 mice per strain total from >3 independent experiments. **c,** Data are shown as mean plus SEM.

Optimal MHCII presentation also requires appropriate stabilization and trafficking of MHCII by the chaperone protein invariant chain^41–43^. We evaluated invariant chain expression in naïve mouse AT2s by flow cytometry and, consistent with prior reports^22, 38^, found that AT2s express intracellular invariant chain (Fig 2b).

Classical MHCII processing also depends on the protein H2-M, which catalyzes the removal of the invariant chain peptide remnant CLIP from the MHCII peptide-binding groove in the late endosome and facilitates the loading of higher affinity peptides^41–43^. We evaluated H2-M expression by qPCR, RNA-sequencing analysis, and flow cytometry. In mice, H2-M is a heterodimeric protein composed of one α chain pairing with one of two β chains, β1 or β2^47^; H2-Mαβ1 and H2-Mαβ2 are thought to function similarly^48, 49^. By qPCR, AT2s expressed detectable levels of transcripts encoding the α and β chains of H2-M (Fig 2c), using β primers that capture both β1 and β2. By flow cytometry AT2s did not express H2-Mαβ2 protein (Fig 2d), suggesting that the predominant isoform of H2-M in naïve mouse AT2s is therefore H2-Mαβ1, which was confirmed by RNA-sequencing analysis (Fig S5a). We evaluated H2-M protein function in AT2s by flow cytometric detection of the CLIP/MHCII complex CLIP/I-A^b^ in B6 mice. If H2-M is functionally present, then the number of CLIP/I-A^b^ complexes will be low, as CLIP peptide will be edited out. AT2s from wild-type mice have low expression of CLIP/I-A^b^ complexes, similar to hematopoietic MHCII^+^ lung cells, and in contrast to both cell types from H2-M-deficient mice that have high levels of CLIP/I-A^b^ (Fig 2e). Furthermore, the peptide editing activity of H2-M can be antagonized by the protein H2-O^50^. We found that H2-O transcript and protein levels are undetectable in naïve AT2s (Fig S5a,d). Altogether, these data demonstrate that AT2s express functional H2-M (H2-Mαβ1) protein.

We also assessed the expression of invariant chain protein and the human H2-M-equivalent, HLA-DM, in human AT2s, by flow cytometry. As in mice, healthy human AT2s expressed both the invariant chain and HLA-DM (Fig S2b).

Besides extracellular antigens, MHCII also presents peptides derived from intracellular proteins^51–53^. For example, in influenza (flu) virus infections in B6 mice, endogenous MHCII presentation of peptides from viral proteins synthesized within infected cells, instead of engulfed extracellular viral material, drives the majority of the antiviral CD4^+^ T cell response^54^. Endogenous MHCII presentation involves a heterogenous network of machinery that are less well characterized compared to the classical pathway, precluding an exhaustive study of specific mediators here. However, one prerequisite for endogenous processing is expression of intracellular antigen by the APC. We assessed whether AT2s produce viral proteins intracellularly after exposure to live influenza virus and found that AT2s expressed both hemagglutinin (HA) as well as nucleoprotein (NP) to high levels (Fig S4). This suggests that AT2s possess abundant substrate material for endogenous processing during flu infections, and this presumably extends to other respiratory viral infections for which they are also main targets.

Beyond peptide/MHCII complex formation, another component of APC function is the provision of costimulation. In general, priming of naïve T cells requires APC expression of the proteins CD80 or CD86, while the reactivation of effector T cells is less costimulation-dependent and may be influenced by a wider spectrum of costimulatory molecules^55, 56^. Prior studies indicate that naïve AT2s do not express CD80 or CD86, but that they express the noncanonical costimulatory molecule ICAM-1^19–21^. We also assessed AT2s for CD80, CD86, and ICAM-1 expression and found that AT2s expressed ICAM-1, but not CD80 or CD86, both at homeostasis and after influenza infection (Fig S5e,f).

Taken together, these data indicate that AT2s possess the requisite characteristics and intracellular machinery to endow them with MHCII antigen presentation capacity; in combination with the expression of ICAM-1 but not CD80/CD86, this would predict that AT2s have the ability to act as APCs, activating effector, but not naïve, CD4^+^ T cells that enter the lung.

### AT2 MHCII contributes to improved respiratory viral disease outcomes

As AT2s express both MHCII and the conventional MHCII-associated APC machinery, we predicted that AT2 MHCII presentation would contribute to immune responses *in vivo*. To test this, we generated mice with an AT2-specific deletion of MHCII by breeding mice to have both (1) a tamoxifen-inducible Cre enzyme controlled by the AT2-specific surfactant protein C (SPC) promoter^57^ and (2) LoxP sites flanking exon 1 of the I-A^b^ β chain on both alleles^58^ (SPC-Cre-ERT2^+/-^ × H2-Ab1^fl/fl^; aka “SPC^ΔAb1^”). Upon tamoxifen administration, SPC^ΔAb1^ mice exhibit uniform deletion of MHCII from AT2s (Fig 3a-c), in contrast to the parental strains whose AT2s retain MHCII: those expressing the Cre enzyme with a wild-type I-A^b^ locus (SPC-Cre-ERT2^+/-^ only; aka “SPC^Cre^”), as well as those with homozygous floxed I-A^b^ alleles but no Cre (H2-Ab1^fl/fl^ only; aka “Ab1^fl/fl^”).

**Figure 3:**
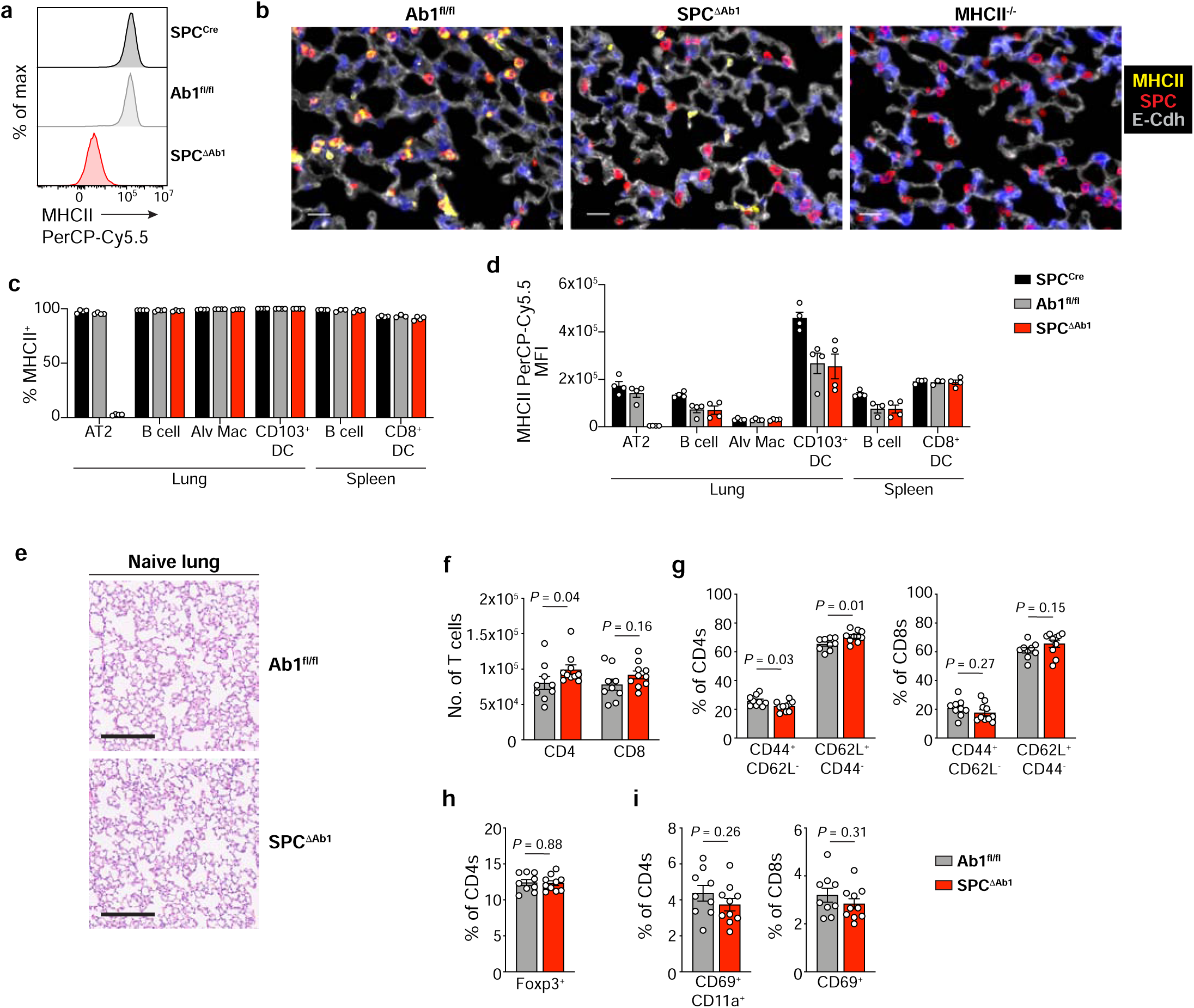
AT2 MHCII is dispensable for immune homeostasis in the lung in 12 week old mice. **a**, MHCII expression by AT2s from naïve SPC-Cre-ERT2^+/-^ (SPC^Cre^), H2-Ab1^fl/fl^ (Ab1^fl/fl^), and SPC-Cre-ERT2^+/-^ × H2-Ab1^fl/fl^ (SPC^ΔAb1^), lungs measured by flow cytometry; histograms represent n>50 mice per strain total from >10 independent experiments. **b**, Immunofluorescence detection of MHCII, SPC, and E-Cadherin in Ab1^fl/fl^, SPC^ΔAb1^, and MHCII^-/-^ lungs; scale bar depicts 20 μm, representative images of n=1-2 mice per strain. **c,d,** Percent and MFI of each cell type expressing MHCII protein for lung AT2s, B cells, alveolar macrophages, CD103^+^ DCs, and spleen B cells and CD8^+^ DCs, from naïve SPC^Cre^, Ab1^fl/fl^, and SPC^ΔAb1^ mice, measured by *ex vivo* flow cytometry analysis; bars show n=4 mice per strain from 1 experiment, representative of 2 similar independent experiments for the Ab1^fl/fl^ and SPC^ΔAb1^ strains (n=8 mice total per strain). **e,** H&E staining of naïve 12 week old Ab1^fl/fl^ and SPC^ΔAb1^ lungs, images representative of n=5 mice per strain. Scale bar depicts 200 μm. **f-i,** Absolute number of lung CD4^+^ and CD8^+^ T cells (**f**), and frequency of lung CD4^+^ and CD8^+^ T cells expressing CD44, CD62L (**g**), Foxp3 (**h**), and CD69, CD11a (**i**), as indicated, from naïve 12 week old Ab1^fl/fl^ and SPC^ΔAb1^ mice measured by *ex vivo* flow cytometry analysis; n=9-10 mice per strain from 1 experiment, representative of 2 independent experiments. **c-d, f-i**, Data shown are mean plus SEM, analyzed by Mann-Whitney test (**f** [CD4s]) or unpaired two-tailed Student’s *t*-test (**f** [CD8s], **g-i**). Full statistical test results are in Extended Data Table 1.

To ensure that the deletion of MHCII in SPC^ΔAb1^ mice was restricted to AT2s, we compared MHCII on various other APCs in the lungs and spleens of all three strains. Similar frequencies of lung B cells, alveolar macrophages, lung CD103^+^ DCs, splenic B cells, and splenic CD8α^+^ DCs, expressed MHCII in all three strains, suggesting that the Cre-mediated deletion of MHCII was AT2-specific (Fig 3c). However, the overall magnitude of MHCII on certain APCs was lower in both the SPC^ΔAb1^ and Ab1^fl/fl^ mice compared to the SPC^Cre^ strain (Fig 3d); this suggests that in mice of the H2-Ab1^fl/fl^ genetic background, the I-A^b^ targeting construct diminishes MHCII expression in some cell types, independent of Cre expression. Based on these data, we considered the Ab1^fl/fl^ mice the most appropriate comparators for the SPC^ΔAb1^ mice, so these two strains were paired for our *in vivo* studies.

We first assessed whether loss of MHCII on AT2s resulted in lung disease or immune dysfunction at homeostasis. To do so, we compared hematoxylin and eosin (H&E) staining of naïve 12-week-old SPC^ΔAb1^ and Ab1^fl/fl^ lungs. Both strains demonstrated normal, healthy lung architecture with minimal infiltrates (Fig 3e). We also examined whether SPC^ΔAb1^ mice had subclinical alterations in lung immune cells as well as splenic T cells, by flow cytometry. 12-week-old old naïve SPC^ΔAb1^ and Ab1^fl/fl^ lungs and spleens had overall similar frequencies and numbers of CD4^+^ and CD8^+^ T cells (Fig 3f, S6, Extended Data Table 1); both T cell subsets in both organs of the two strains also demonstrated similar expression of memory phenotype markers CD44 and CD62L (Fig 3g, S6), the proliferation marker Ki-67, and the inhibitory receptor and activation/exhaustion marker PD1 (Fig S6). Minor differences between SPC^ΔAb1^ and Ab1^fl/fl^ mice achieved statistical significance in the cases of CD4^+^ T cell numbers, the proportion of CD4s expressing CD44/CD62L, and the proportions of CD4s and CD8s expressing the activation/exhaustion marker PD1. However, these differences do not result in detectable pathology, as above (Fig 3e), nor do they appear to influence outcomes following infection, as described below (Fig S9). As surface MHCII is the main ligand for the T cell inhibitory receptor LAG3, which restricts T cell expansion and effector function^59^, we also evaluated LAG3-expressing T cells in the lungs and spleen of both strains at homeostasis; we found no differences in the frequencies of LAG3^+^ CD4^+^ and CD8^+^ T cells (Fig S7, Extended Data Table 2). The frequency of lung and splenic Tregs was also similar between SPC^ΔAb1^ and Ab1^fl/fl^ mice (Fig 3h, S6). Frequencies of tissue resident-memory CD69^+^CD11a^+^ CD4^+^ and CD69^+^ CD8^+^ T cells^60^ in the lungs were also similar between the two strains (Fig 3i), as were the frequencies and numbers of alveolar macrophages, neutrophils, B cells, NK cells, and γδT cells (Fig S6). Thus, overall, MHCII appears to be dispensable for the maintenance of a healthy immune environment in the lung at homeostasis in young adult mice. Furthermore, as the SPC^ΔAb1^ mice did not experience respiratory disease at homeostasis, and AT2s sorted from SPC^ΔAb1^ and Ab1^fl/fl^ mice formed similar numbers of lung organoids *in vitro* (Fig S8), MHCII also does not seem to be required for two main physiologic functions of AT2s: surfactant production and lung regeneration.

We next assessed whether AT2 MHCII contributes to the outcome of lung infection. To do this, we used two respiratory virus infection models: mouse lung-adapted PR8 influenza A virus (IAV) as well as Sendai virus (SeV), a natural mouse pathogen similar to human parainfluenza virus. To assess the impact of AT2 MHCII on viral disease, we measured weight loss of SPC^ΔAb1^ and Ab1^fl/fl^ mice after infection with either IAV or SeV. SPC^ΔAb1^ and Ab1^fl/fl^ mice exhibited similar weight loss after IAV infection (Fig 4a; Extended Data Table 3). However, after SeV infection, SPC^ΔAb1^ mice experienced significantly more weight loss and delayed recovery compared to the Ab1^fl/fl^ group (Fig 4b; Extended Data Table 4). We also assessed the impact of AT2 MHCII on mortality after infection with both viruses. SPC^ΔAb1^ mice experienced reduced survival after IAV infection compared to Ab1^fl/fl^ controls, 24% vs 50% respectively (Fig 4c). Similarly, after SeV infection, SPC^ΔAb1^ mice had worse survival with approximately 2-fold higher mortality (28%) compared to the Ab1^fl/fl^ controls (12%) (Fig 4d). We next asked whether AT2 MHCII antigen presentation might contribute to the control of viral replication, by measuring lung virus titers after infection with either IAV or SeV. There were no significant differences in IAV titers 7 days after infection between SPC^ΔAb1^ and Ab1^fl/fl^ mice (Fig 4e). Similarly, SeV titers 4, 7, and 9 days after infection were similar between the two strains (Fig 4f). As LAG3 has been shown to restrict lung CD4^+^ and CD8^+^ T cell responses during respiratory viral infections^61, 62^, we also considered whether AT2 MHCII is required for LAG3-mediated T cell suppression during infection. We found similar frequencies and numbers of LAG3^+^ CD4^+^ and CD8^+^ T cells in the lungs of SPC^ΔAb1^ and Ab1^fl/fl^ mice 9 days after IAV infection (Fig S7). Thus, while MHCII on AT2s is not required for protection against respiratory viral infection, it does contribute to lower morbidity and mortality, without impacting LAG3^+^ T cell expansion or lung viral burden.

**Figure 4:**
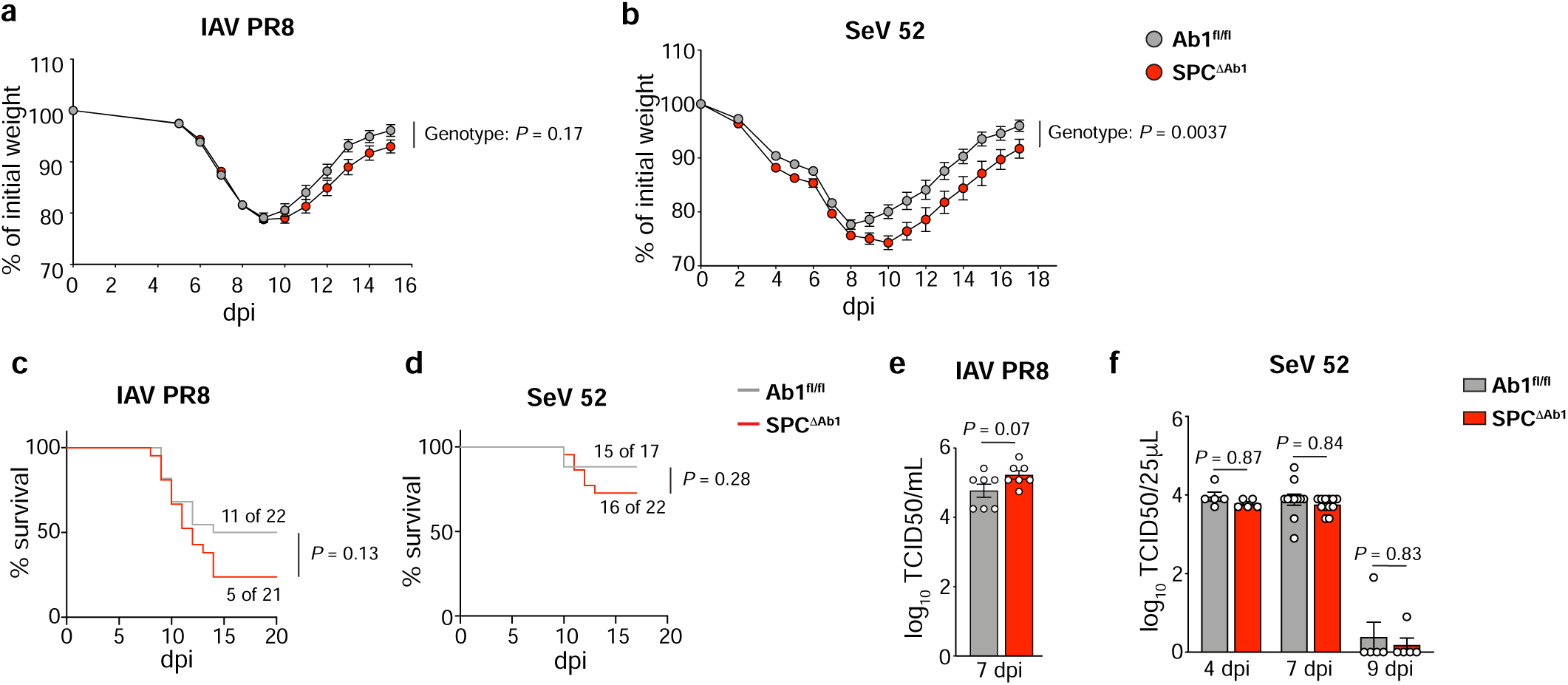
Loss of AT2 MHCII results in greater weight loss and reduced survival after respiratory viral infection. **a-f**, Comparison of Ab1^fl/fl^ and SPC^ΔAb1^ mice after infection with influenza strain PR8 (IAV PR8) (**a,c,e**) and Sendai virus strain 52 (SeV 52) (**b,d,f**). **a**,**b,** Weight loss with weights displayed relative to day of infection; n=43-53 mice per strain pooled from 6 independent experiments (**a**), and n=17-22 mice per strain pooled from 2 independent experiments (**b**). **c,d**, Mortality, with curves representing proportion surviving; n=21-22 mice per strain pooled from 2 independent experiments (**c**), and n=17-22 mice per strain pooled from 2 independent experiments (**d**). **e, f,** Lung virus titers on the days post-infection (dpi) indicated; n=7 mice per strain from 1 experiment (**e**), and n=5 mice per strain for days 4 and 9, n=11-12 per strain for day 7, pooled from 2 independent experiments (**f**). **a,b**, Data are mean plus SEM, analyzed by mixed-effects model with post-hoc multiple comparisons with Sidak’s correction at each dpi. *P* values displayed represent overall model effects. Full statistical test results are in Extended Data Tables 3-4. **c,d**, Survival curves were compared via Log-rank test (χ^2^=2.327, df=1 (**c**), and χ^2^=1.173, df=1 (**d**)). **e,f,** Bars are mean plus SEM of log_10_ transformed values, analyzed by unpaired two-tailed Student’s *t*-test (*t*=1.995, df=12) (**e**), and two-way ANOVA (genotype effect: *F*=1.559, df.n=1, df.d=37, *P*=0.2197) with post-hoc multiple comparisons with Sidak’s correction (4 dpi *t*=0.6853, df=37; 7 dpi *t*=0.7603, df=37; 9 dpi *t*=0.7614, df=37). *P* values displayed represent post-hoc comparisons (**f**).

We also asked whether the minor, yet statistically significant, differences between certain lung T cell subsets in SPC^ΔAb1^ and Ab1^fl/fl^ mice observed at homeostasis were amplified during infection. We observed no statistically significant differences in the numbers of lung CD4^+^ T cells or the proportion of lung CD4s expressing CD44/CD62L between SPC^ΔAb1^ and Ab1^fl/fl^ mice 9 days after IAV infection (Fig S9a,b; Extended Data Table 5). Furthermore, the numbers and frequencies of lung CD4 and CD8 T cells expressing the activation/exhaustion marker PD1 were similar (Fig S9c,d), as was the proportion of AT2s expressing PD-L1, the cognate ligand for PD1 (Fig S9e). Thus, the slight differences in magnitude of these T cell subsets at homeostasis are unlikely to explain the differences in outcomes following infection.

In summary, these studies suggest that *in vivo* in young adult mice, AT2 MHCII expression is dispensable for healthy lung immune homeostasis but confers an appreciable advantage in respiratory viral disease outcome overall.

### AT2s exhibit restrained antigen presentation capacity via MHCII

Although loss of AT2 MHCII worsened respiratory virus disease, the effect was smaller than anticipated based on the abundance of AT2s in the lung^32^ and the magnitude of AT2 MHCII expression. A potential explanation for this more measured impact is that AT2s possess limited MHCII antigen processing and presentation capacity.

To investigate the antigen presentation function of AT2s, we first evaluated the ability of AT2s to stimulate flu peptide/MHCII complex-specific costimulation-independent T cell hybridomas (Fig 5a). AT2 presentation of five different epitopes from live virus was undetectable (Fig 5b,c), in contrast to professional APCs that presented all five. Poor presentation by AT2s was not due to a failure of *in vitro* infection, as AT2s were infected to levels higher than comparator professional APCs (Fig S4a). Limited presentation was observed across multiple MHCII alleles, B6 I-A^b^ (Fig 5b) and BALB/c I-E^d^ (Fig 5c), was not the result of protein source, as both neuraminidase (NA) and HA-derived epitopes were similarly affected (Fig 5b, c), and was similar for epitopes generated by both exogenous (HA^107–119^) and endogenous (NA^79–93^) processing pathways^63^ (Fig 5c). Even when pulsed with synthetic peptides, AT2s were able to present only three out of five epitopes (HA^91–107^, HA^302–313^, NA^79–93^) (Fig 5b,c). Thus, AT2s exhibited a global impairment in the capacity to present MHCII-restricted epitopes *in vitro*.

**Figure 5:**
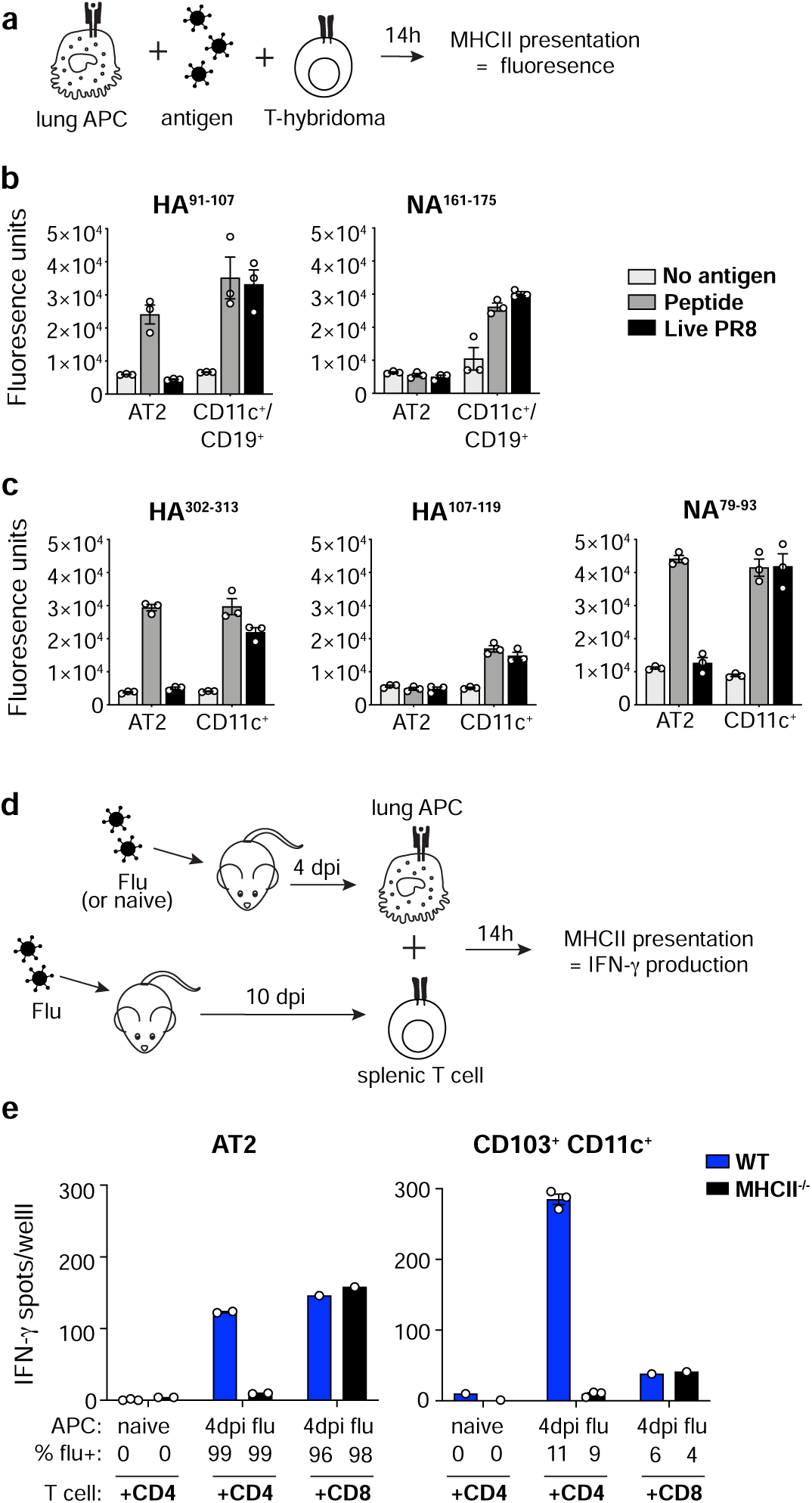
AT2s exhibit a globally restricted capacity to present influenza virus epitopes via MHCII. **a,** Schematic of hybridoma presentation assay. **b**,**c**, Presentation of MHCII-restricted flu peptides by B6 AT2s and a mixed population of CD11c^+^ and CD19^+^ lung cells (**b**) or BALB/c AT2s and CD11c^+^ lung cells (**c**) sorted from naïve mice then incubated with synthetic peptide or live virus, reflected by NFAT-*LacZ-*inducible T cell hybridoma activation and cleavage of a fluorogenic β-galactosidase substrate; bars shown represent 3 technical replicates plus SEM from 1 experiment, representative of 3 similar independent experiments for (**b**). **d,** Schematic of primary T cell ELISpot presentation assay. **e,** Presentation of MHCII-restricted flu peptides by lung AT2s and CD103^+^ CD11c^+^ APCs sorted from naïve or flu-infected WT or MHCII^-/-^ B6 mouse lungs 4 dpi, measured as the production of IFN-γ by responding flu-experienced splenic CD4^+^ and CD8^+^ T cells (as indicated); “APC %flu+” numbers describe the proportion of each APC type that was flu-infected (shown in Fig S4b), and bars shown represent 1-3 technical replicates plus SEM from 1 experiment, representative of 2-3 similar independent experiments.

Assays of AT2 function *in vitro* may be suboptimal due to reduced viability after cell sorting and the gradual de-differentiation of AT2s in standard tissue culture^64, 65^. Additionally, measuring presentation on an individual epitope basis may underestimate the presentation of all possible MHCII-restricted flu peptides. Thus, we next examined AT2 *in vivo* flu peptide/MHCII complex formation, by co-culturing *in vivo*-infected AT2s from wild-type or MHCII^-/-^ B6 mice taken 4 days post influenza infection with polyclonal splenic CD4^+^ or CD8^+^ T cells taken 9 days post-infection, in the presence of soluble anti-CD28 (Fig 5d). MHCII presentation was detected via T cell IFNγ production captured by ELISpot. AT2s from flu-infected mice were capable of stimulating both CD4^+^ and CD8^+^ T cells to produce IFNγ (Fig 5e). Stimulation of CD4^+^ T cells, but not CD8^+^ T cells, was abrogated when MHCII^-/-^ flu-infected AT2s were used as APCs, confirming that stimulation of CD4^+^ T cells by AT2s was MHCII-dependent. However, consistent with our *in vitro* studies, AT2 presentation to CD4^+^ T cells was markedly less efficient than professional APCs, in this case CD103^+^ CD11c^+^ cells taken from the same lungs; despite being 9 times more infected, AT2s (99% infected, Fig S4b) stimulated less than half the IFNγ production than the comparator APCs (11% infected, Fig S4b).

We considered that the high degree of AT2 infection in our assays might impair their capacity to process and present antigen. To address this, we measured MHCII antigen presentation using a non-infectious model system, by staining lungs directly *ex vivo* with the peptide/MHCII complex-specific “YAe” antibody^66^ (Fig 6a). YAe detects Eα^52–68^/I-A^b^ complexes, which form in [BALB/c x B6] F1 mice as they are composed of a peptide from I-E^d^ (BALB/c-derived) presented by I-A^b^ (B6-derived). In naïve F1 mice, AT2s had low yet detectable YAe staining above the background levels of B6 AT2s (Fig 6b,c). However, YAe staining of F1 AT2s was significantly lower than that of F1 lung B cells (Fig 6b,c, Extended Data Table 6). This could not be explained by differences in expression of the source proteins, I-A^b^ and I-E^d^, which are similarly expressed by AT2s and B cells at homeostasis, if not slightly higher in AT2s (Fig 6d,e, Extended Data Tables 7-8). These results indicate that AT2s are capable of forming Eα^52-68^/I-A^b^ complexes at steady state, but they do so far less efficiently than do B cells.

**Figure 6:**
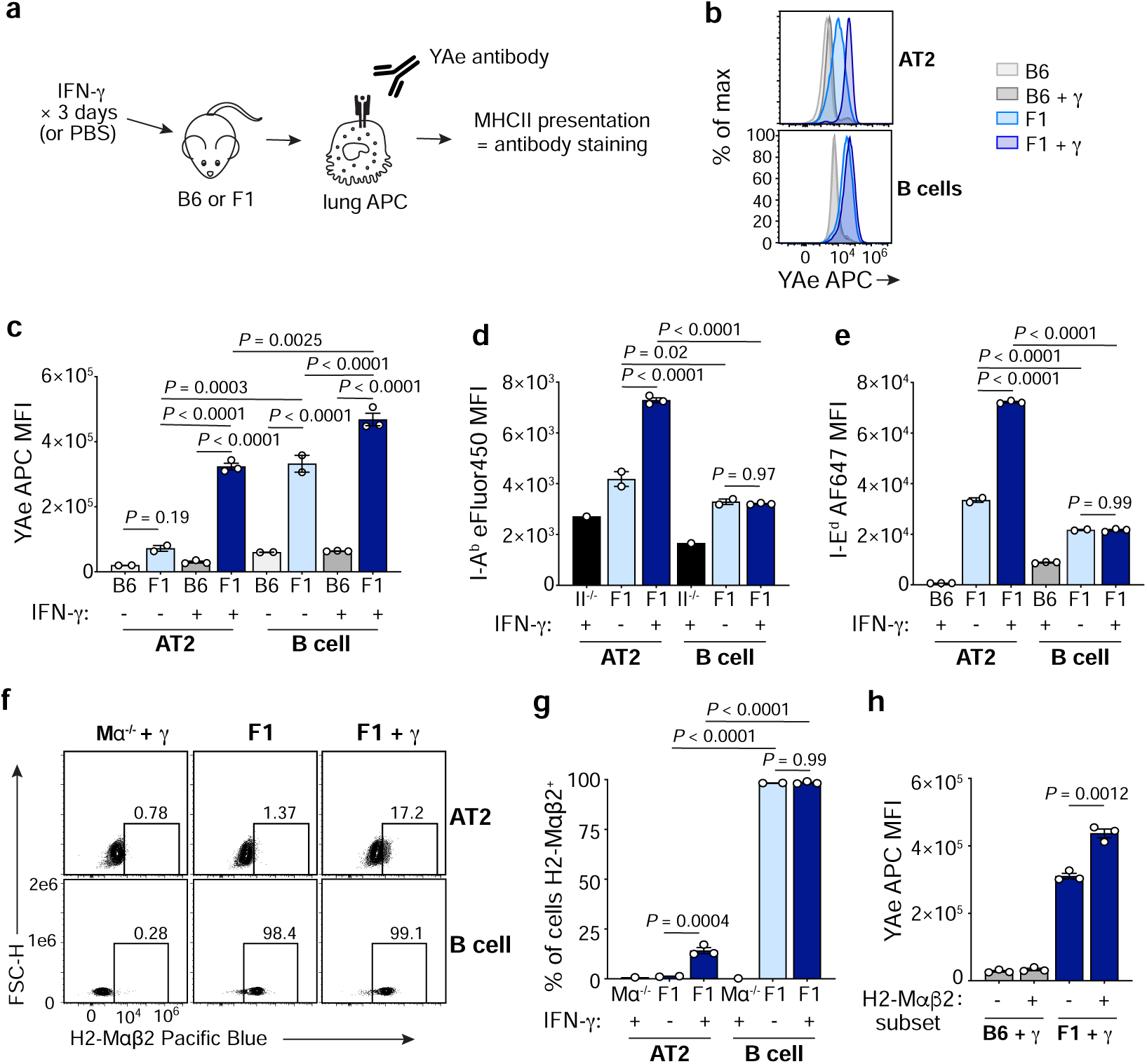
AT2 MHCII presentation is enhanced in the setting of inflammation but remains limited. **a,** Schematic of YAe presentation assay. **b,c,** Eα^52–68^/I-A^b^ complex formation by AT2s and B cells, detected by *ex vivo* YAe antibody staining of lungs from WT B6 or F1 [BALB/c x B6] mice treated with PBS or IFNγ. **d,e,** I-A^b^ (**d**) and I-E^d^ (**e**) expression by AT2s and B cells, detected by *ex vivo* flow cytometry staining of lungs from MHCII^-/-^ (II^-/-^) B6 (**d**), WT B6 (**e**), or F1 (**d,e**) mice treated with PBS or IFNγ (as indicated). **f,g,** Intracellular H2-Mαβ2 expression by AT2s and lung B cells from F1 or *H2-DMα*^-/-^ (Mα^-/-^) B6 mice treated with PBS or IFNγ. Frequency of H2-Mαβ2^+^ cells is shown above gates (**f**), which were drawn separately for AT2s and B cells based on corresponding *H2-DMα*^-/-^ cells. **h,** YAe staining of H2-Mαβ2^+^ and H2-Mαβ2^-^ subpopulations of AT2s from B6 and F1 mice treated with IFNγ. **b, f,** Plots represent n=4-6 mice per group total from 2 independent experiments. **c-e,g,h,** Bars represent mean plus SEM and depict n=2-3 mice per group from 1 experiment, representative of 2 similar independent experiments (total n=4-6) (**c,g,h**) or n=1-3 mice per group from 1 experiment (**d,e**). Data were analyzed by three-way ANOVA with multiple comparisons with Sidak’s correction (**c**), two-way ANOVA with multiple comparisons with Tukey’s correction (**d,e,g**), and unpaired two-tailed Student’s *t*-test (*t*=8.165, df=4) (**h**). *P* values displayed represent post-hoc comparisons of interest (**c-e,g**). Full statistical test results are in Extended Data Tables 6-9.

To better approximate the setting of viral infection, we also treated mice with IFNγ and then measured Eα^52-68^/I-A^b^ complex formation. AT2 YAe staining was significantly increased in F1 mice after treatment with IFNγ, but was again lower than B cells in the same mice (Fig 6b,c). To assess the contributors to increased MHCII presentation in AT2s, we evaluated changes in expression of I-A^b^, I-E^d^, and H2-M, all of which are required for Eα^52-68^/I-A^b^ complex formation^67^. Both I-A^b^ and I-E^d^ increased in AT2s after IFNγ administration (Fig 6d,e). IFNγ also induced some AT2s to express the H2-Mαβ2 isoform of H2-M (Fig 6f,g, Extended Data Table 9), which is absent at baseline where AT2s express the H2-Mαβ1 isoform only, as previously demonstrated (Fig 2, S5a). Although H2-Mαβ2^+^ AT2s exhibited more YAe staining than H2-Mαβ2^-^ AT2s, both populations stained highly after IFNγ treatment (Fig 6h), suggesting that a higher magnitude of H2-M contributes to enhanced Eα^52-68^/I-A^b^ presentation, but the specific H2-Mαβ2 isoform is not required. Thus, AT2 MHCII presentation increases during IFNγ inflammation due to the upregulation of several MHCII presentation mediators, but it is still less efficient than that of professional APCs.

Taken together, these data demonstrate that AT2s are capable of presenting antigen via MHCII to CD4^+^ T cells, and this is enhanced in the setting of inflammation; however, compared to professional APCs they exhibit limited antigen presentation of MHCII-restricted epitopes derived from both viral and endogenous proteins, as well as some extracellular peptides. This restrained presentation capacity may explain the variability seen in prior studies of AT2 function owing to differences in the model systems used, in particular the level of antigen provided to the AT2s, as well as the specific epitopes studied.

## Discussion

Here we establish that AT2 cells stand alone in a new category of MHCII-expressing cells, as non-immune cells that nonetheless constitutively express MHCII in an IFNγ-independent manner. We demonstrate that AT2 MHCII may contribute to improved outcomes of respiratory virus disease *in vivo,* and that in contrast to professional APCs, AT2s exhibit a program of restrained MHCII antigen presentation.

We were unable to identify an immune signal required for AT2 MHCII expression despite investigating a variety of broad adaptive and innate mediators, including the microbiota. This suggests that MHCII expression is not driven by inflammation. As AT2 MHCII is upregulated only after birth (J. Whitsett, B. Aronow, S. Potter [CCHMC] and the LungMAP Consortium [U01HL122638]; single cell RNA-seq data available at www.lungmap.net), its expression may instead be induced by a non-immunologic environmental stimulus or may be an intrinsic part of the AT2 cell developmental identity. The constitutive nature of AT2 MHCII expression prompted us to also consider whether AT2 MHCII plays a novel, non-immune role in basic AT2 biology. We demonstrated that AT2 MHCII is not required for the regenerative function of AT2s, and as the SPC^ΔAb1^ mice exhibit no lung disease either clinically or histologically, it is most likely also dispensable for surfactant production. More extensive characterization of AT2 physiology and function will be needed to investigate whether there are other non-immunologic processes that are disrupted by the loss of MHCII from AT2s.

We demonstrated that inefficient antigen presentation by AT2s cannot be explained by the absence of conventional MHCII-associated machinery. Instead, it is possible that there are mechanisms operating in AT2s that actively oppose MHCII presentation. AT2s possess a highly specialized endosomal network specifically tailored to the synthesis, secretion, and recycling of surfactant proteins and lipids^46^. It is possible that these endocytic compartments are incompatible with the spatiotemporal degradative and peptide loading requirements for MHCII presentation. It is also unusual that AT2s present only some synthetic peptides when provided exogenously. One possible explanation is that AT2 cell surface MHCII molecules are already bound to a high affinity self-peptide that can be displaced only by even higher affinity peptides.

Restrained MHCII antigen presentation by AT2s is consistent with the more measured contribution they make via MHCII to the outcome of lung viral infection. We propose that this limited presentation is favorable in the lung, where excessive T cell activation would be especially damaging. In this model, high MHCII expression poises AT2s to amplify lung adaptive immune responses, but restricted MHCII presentation establishes a higher threshold that must be overcome in order to do so. This system would allow for a more tempered response, where MHCII presentation by AT2s would be sufficient to trigger cognate T cell activation only in the setting of high antigen burden, such as a severe lung infection, but would prevent AT2-induced amplification of T cells in response to low levels of inhaled innocuous environmental antigens. Ongoing and future work will further explore this possibility.

We found that AT2 MHCII did not affect the total amount of virus in the lung, suggesting that AT2 MHCII contributes to protection by other mechanisms. AT2 MHCII may instead facilitate the development of a protective CD4^+^ T cell cytokine milieu, which is critical for disease outcome in the setting of respiratory viral infection^68, 69^. CD4^+^ Tregs have also been demonstrated to facilitate lung repair by enhancing AT2 proliferation^70, 71^. Thus, it is also possible that AT2 MHCII helps to facilitate disease recovery by promoting the expansion or localization of CD4^+^ Tregs to the regions of the lung experiencing high virus loads, in order to minimize damage and enhance alveolar regeneration.

As lung-resident memory CD4^+^ (Trm) can mediate protection upon viral challenge^72^ and circulating virus-specific memory CD4^+^ T cells correlate with heterosubtypic protection in humans^73^, in future studies it will be important to address whether AT2 MHCII contributes to the development of virus-specific memory CD4^+^ T cells. Prior studies from the Swain laboratory suggest that robust influenza-specific memory CD4^+^ T cell formation requires two cognate TCR-MHCII encounters: the first priming interaction in the lymph node, and a second around days 5-7 post-infection^74, 75^. As MHCII^+^ AT2s are present at the site of infection with access to both antigen and antigen-specific CD4^+^ T cells, it is possible that AT2s help to deliver this second checkpoint. It is also possible that local AT2 MHCII antigen presentation helps to facilitate the development and retention of lung-resident virus-specific memory CD4^+^ Trm during infection^76–78^.

AT2 MHCII may also play an important role in regulating CD4^+^ T cell-driven lung diseases in response to respiratory allergens, such as in allergic asthma as well as hypersensitivity pneumonitis^79–82^. We demonstrated that the loss of AT2 MHCII did not result in lung disease at homeostasis in 12-week-old specific pathogen free (SPF) mice, suggesting that AT2 MHCII does not contribute to inhaled antigen tolerance or allergy. However, as SPF mice experience artificially low levels of antigen exposure, it is possible that the impact of AT2 MHCII loss would be more apparent in mice with a greater burden or longer duration of inhaled putative allergen exposure, such as aged mice or wild mice^83–86^, or those intentionally sensitized to allergens.

Finally, an essential avenue of future work will also be to understand whether alterations in AT2 MHCII expression or function contribute to the wide variation in outcome of infectious and immunologic lung diseases in humans. Of special interest at this time is whether AT2 MHCII contributes to protection in the setting of SARS-CoV-2 infection, as AT2s are the main cell type infected with this virus in the human lung parenchyma^87–89^.

## Methods

### Mice

C57Bl/6 wild-type (B6), B6.129S2-*H2^dlAb1-Ea^*/J (MHCII^-/-^)^90^, B6;129S4-*H2-DMa^tm1Luc^*/J (*H2-DMa*^-/-^)^91^, B6.129S7-*Ifng^tm1Ts^*/J (*Ifng*^-/-^)^92^, B6.129S(Cg)-*Stat1^tm1Dlv^*/J (*Stat1*^-/-^)^93^, C.129S2(B6)-*Ciita^tm1Ccum^*/J (*Ciita*^-/-^)^94, 95^, B6.SJL-*Ptprc^a^ Pepc^b^*/BoyJ (CD45.1 B6), B6.129X1-*H2-Ab1^tm1Koni^*/J (*H2-Ab1*^fl/fl^)^58^, as well as BALB/c, and CB6F1/J (F1 [BALB/c x C57Bl/6] mice were originally purchased from the Jackson Laboratory. C57Bl/6 *Cd74*^-/-^ mice^96^ were originally provided by Guo-Ping Shi (Harvard), C57Bl/6 *Ciita pIV*^-/-^ mice^39^ were provided by S. Hugues (University of Geneva), and C57Bl/6 *H2-Ob^-/-^* mice^97^ were provided by L. Denzin (Rutgers). C57Bl/6 SPC-Cre-ERT2 mice^57^ were provided by G.S. Worthen, B6.129S7-*Ifngr1^tm1Ag^*^t^/J (*Ifngr1*^-/-^)^98^, B6(Cg)-*Ifnar1^tm1.2Ees^*/J (*Ifnar1*^-/-^)^99^, and B6.Cg-*Ifngr1^tm1Agt^ Ifnar1^tm1.2Ees^*/J (*Ifnar1*^-/-^*Ifngr1*^-/-^) mice were provided by E. Behrens, and germ-free mice were provided by M. Silverman (Children’s Hospital of Philadelphia). C57Bl/6 B6.129P2(SJL)-*Myd88^tm1.1Defr^*/J (*Myd88*^-/-^) mice^100^ were provided by S. Shin, and B6.129S2(C)-*Stat6^tm1Gru^*/J (*Stat6*^-/-^) mice^101^ were provided by C. Hunter (University of Pennsylvania). All mice were maintained in specific pathogen-free facilities at the Children’s Hospital of Philadelphia, except for *Myd88*^-/-^ and *Stat6*^-/-^ mice, which were housed in specific pathogen-free facilities at the University of Pennsylvania, germ-free mice, which were housed in gnotobiotic mouse facilities at the University of Pennsylvania, and *H2-Ob^-/-^* mice, which were housed in the animal facility at Rutgers University. Mice were age and sex matched for all studies and 6-12 week old mice were used for all experiments, unless otherwise specified. All animal procedures were in compliance with institutional and AAALAC ethical guidelines and were approved by the Institutional Animal Care and Use Committee (IACUC) at the Children’s Hospital of Philadelphia.

### Cell lines

The B6 primary skin fibroblast cell line was derived in our laboratory and has been described previously^102^; it was maintained in Dulbecco’s Modified Eagle Medium (DMEM) containing 5% FBS, Penicillin-Streptomycin, and 2mM L-glutamine. Madin-Darby canine kidney (MDCK) cells were provided by S. Hensley (University of Pennsylvania) and were maintained in MEM with 10% FBS. Monkey kidney LLC-MK2 cells were provided by Carolina Lopez (University of Pennsylvania) and were maintained in DMEM with 10% heat-inactivated FBS, Penicillin-Streptomycin, 2 mM L-glutamine, and 1 mM sodium pyruvate. T cell hybridoma lines were derived in our laboratory and have been described previously^54, 63^; these were maintained in “R10 media”: RPMI media containing 10% FBS, 50 μM 2-mercaptoethanol, Penicillin-Streptomycin, 2mM L-glutamine.

### Viruses

Mouse lung-adapted H1N1 influenza A virus, A/Puerto Rico/8/1934 (IAV PR8), was originally provided by C. Lopez (University of Pennsylvania), and then expanded in 10-day-old embryonated chicken eggs. IAV PR8 viral titer was determined by standard focus-forming unit (FFU) assay in MDCK cells. Sendai virus, strain 52 (SeV 52), was also a generous gift of C. Lopez (University of Pennsylvania). SeV 52 viral titer was determined using a tissue culture infectious dose (TCID_50_) standard infectivity assay in LLC-MK2 cells.

### Synthetic peptides

The following synthetic peptides were used: HA^91–107^ (RSWSYIVETPNSENGIC), NA^161–175^ (SVAWSASACHDGMGW), HA^302–313^ (CPKYVRSAKLRM), HA^107–119^ (SVSSFERFEIFPK), NA^79–93^ (IRGWAIYSKDNSIRI). All peptides were obtained at >85% purity from Genscript.

### Tissue isolation for *in vitro* analysis and flow cytometry

For isolation of AT2 cells, the lung vasculature was first perfused with 5 mL PBS by injecting into the cardiac right ventricle. Lungs were then inflated via intratracheal instillation of 0.9 mL AT2 digest media [5 U/mL Dispase (354235, Corning), 1.66 mg/mL Collagenase A (10103586001, Sigma), 0.33 mg/mL DNase I (10104159001, Sigma) in PBS]. The trachea was then tied with suture to keep the lungs inflated while they were excised en bloc and then placed in an additional 1 mL digest media, then incubated at 37°C. After 45 minutes, 5 mL 20% FBS in PBS was added, and the parenchymal lung lobes were removed from the large airways with forceps, then dissociated by vigorous pipetting. Digested lungs were passed through a 70 μm strainer, incubated in ACK lysis buffer to remove RBCs, then passed through a 40 μm strainer to obtain a single cell suspension.

For isolation of lung T cells, the lung vasculature was first perfused with 5 mL 1% FBS in PBS by injecting into the cardiac right ventricle. Individual parenchymal lung lobes were removed from the chest cavity and placed into gentleMACS C tubes containing 2 mL 1% FBS in PBS. Lymphocyte digest media was then added to the lungs (1% FBS, 2.25 mg/mL Collagenase D (11088866001, Sigma), 0.15 mg/mL DNase I in PBS in 4 mL final volume), which were then gently disrupted using gentleMACS homogenizer program m_spleen_01.01, then incubated for 45 minutes at 37C with shaking. R10 media was then added to each tube, followed by further dissociation using gentleMACS homogenizer program m_lung_02.01. Digested lungs were then passed through a 70 μm strainer, incubated in ACK lysis buffer to remove RBCs, then passed through a 40 μm strainer to obtain a single cell suspension.

AT2s could not be recovered from the T cell digest; likewise, the AT2 digest was not optimal for T cell isolation as it resulted in the degradation of surface CD4 and CD8 from T cells. In some T cell isolation experiments, a small sample of AT2s was also needed for the purposes of confirmation phenotyping to assess deletion or presence of MHCII. In these cases, prior to the addition of lymphocyte digest media to the lungs, a small ([<2mm]^3^) portion of lung was removed, and then incubated for 1 hour at 37C in 0.4 mL AT2 digest media. Next, 0.1 mL of FBS was added, and the sample was then homogenized through a 40 μm cell strainer using the flat end of a 1 mL syringe plunger, to obtain a single cell suspension.

To harvest lungs for virus titering, lung lobes were removed directly from the thoracic cavity (with no perfusion or instillation) and placed in gentleMACS M tubes containing 1 mL PBS. Additional PBS was then added to each M tube to produce a final 10% weight/volume solution of lungs/PBS. Lungs were then homogenized using gentleMACS dissociator program RNA_01.01. Cellular debris was removed by centrifugation at 600xg for 10” at 4°C, and the clarified supernatant was then used for virus titering.

For isolation of splenocytes, spleens were removed from the abdominal cavity and placed directly in PBS. Spleens were then homogenized through a 70 μm cell strainer using the blunt end of a 3 mL syringe plunger. The splenocytes were then incubated in ACK lysis buffer to remove RBCs, then passed through a 40 μm strainer to obtain a single cell suspension.

For isolation of immune cells from peripheral blood, blood was collected via cheek bleed or IVC puncture into PBS containing 25 mM EDTA. Samples were centrifuged at 300xg for 10”, followed by ACK lysis of the cell pellet to remove RBCs.

For isolation of bone marrow cells, whole bones of the hindlimb were removed from mice. In a sterile manner, the ends of the bones were cut and the interior cavity was flushed with PBS.

For isolation of human distal lung cells, healthy human lungs were obtained with informed consent from the Prospective Registry of Outcomes in Patients Electing Lung Transplant Study approved by University of Pennsylvania Institutional Review Board in accordance with institutional ethical procedures and guidelines. The donor used in this study had the following characteristics: 36 y.o. M, no smoking history, no evidence of active infection, P/F ratio of 346. Lungs were digested as described previously by Zacharias and Morrisey^103^. Briefly, a 2×2 cm piece of distal lung tissue (pleura and airways removed) was minced then processed in the same AT2 digest media as above using a gentleMACS dissociator at 37°C for 35 minutes. Digested lungs were washed, passed through 70 μm and 40 μm strainers, and then incubated in ACK lysis buffer to remove RBCs and generate a single cell suspension.

### Cell type identification via flow cytometry and cell sorting

All flow cytometry antibodies used are listed in Supplementary Table 1. For all flow cytometry experiments, unless otherwise stated, single cell suspensions were first stained for dead-cell exclusion with Live/Dead Aqua (L34957, Thermo Fisher) in PBS, followed by Fc-receptor blockade with anti-CD16/CD32 (553141, BD) for mouse cells or Human TruStain FcX (422301, Biolegend) for human cells, in 0.1% BSA in PBS. Cells were then stained with surface antibodies diluted in 0.1% BSA in PBS for 30” in the dark at 4°C. If necessary, cells were then fixed and permeabilized for intracellular staining using the BD Cytofix/Cytoperm kit (554714, BD) or for intranuclear staining using the eBioscience Foxp3 / Transcription Factor Staining kit (00-5523-00, Thermo Fisher). Intracellular and intranuclear stains were diluted in the corresponding kit wash buffer and were incubated for 30” at room temperature. Flow cytometry data were acquired on the following instruments: LSRII and LSRFortessa (BD), CytoFLEX LX and CytoFLEX S (Beckman Coulter). Cell sorting was performed on the following instruments: FACSAria Fusion and FACSJazz (BD), and MoFlo Astrios (Beckman Coulter). All flow cytometry analyses were conducted using FlowJo software (FlowJo LLC).

Murine AT2s were identified by flow cytometry and sorting as detailed in Fig S1^22^. In studies where MHCII expression was evaluated, AT2s were identified using the gating strategy without MHCII as a selection marker, as CD45^-^, CD31^-^, EpCAM^int^ cells. For all other flow cytometry studies, AT2s were identified as CD45^-^, CD31^-^, EpCAM^+^ MHCII^+^ cells. Both gating strategies were validating by intracellular staining for pro-SPC. Human AT2s were identified by flow cytometry as detailed in Fig S2 as HT2-280^+^ lung cells^35^, by staining with unlabeled anti-HT2-280 followed by an anti-mouse IgM fluorophore-conjugated secondary antibody.

Other cell types were identified via flow cytometry as follows: Lung endothelial cells (CD45^-^, CD31^+^), Lung CD103^+^ DCs (CD45^+^, CD11c^+^, CD103^+^), Lung alveolar macrophages (CD45^+^, CD11c^+^, CD103^-^, CD11b^-^, CD64^+^), Splenic CD8^+^ DCs (CD45^+^, CD11c^+^, CD3^-^, CD8^+^), B cells (FSC-A^low^, CD45^+^, CD11c^-^, CD19^+^ or B220^+^), CD4^+^ T cells (CD45^+^, CD3^+^, CD4^+^, CD8^-^), CD8^+^ T cells (CD45^+^, CD3^+^, CD8^+^, CD4^-^), NK cells (CD45^+^, CD3^-^, NK1.1^+^), γδ T cells (CD45^+^, CD3^+^, TCRδ^+^), and neutrophils (CD45^+^, CD11c^-^, CD11b^+^, Ly6G^+^).

For cell sorting (FACS) experiments, samples were not stained with the Live/Dead viability dye or Fc-receptor blockade to minimize sample processing time and preserve cell viability. The following professional APC populations were sorted: qPCR DCs (CD45^+^,CD11c^+^,MHCII^hi^); DQ-ova assay DCs (CD45^+^, CD11c^+^, MHCII^hi^) and B cells (CD45^+^, B220^+^, MHCII^+^); Cathepsin L assay B cells (CD45^+^ CD19^+^); C57Bl/6 hybridoma assay CD11c^+^/CD19^+^ APCs (CD45^+^, MHCII^+^, CD11c^+^ or CD19^+^); Balb/c hybridoma assay CD11c^+^ APCs (CD45^+^, MHCII^+^, CD11c^+^); ELISpot CD103^+^ DCs (CD45^+^, CD11c^+^, CD103^+^). The following AT2 populations were sorted: DQ-ova assay, qPCR, C57Bl/6 hybridoma assay and Balb/c hybridoma assay AT2s (CD45^-^, CD31^-^, EpCAM^+^, MHCII^+^), cathepsin D, cathepsin L, and ELISpot assay AT2s (CD45^-^, CD31^-^, EpCAM^+^), and AT2s for organoid culture (CD45^-^, CD31^-^, Podoplanin^-^, CD34^-^, Sca1^-^, EpCAM^int^). Lung fibroblasts were also sorted for organoid culture as Pdgfrα^+^ cells.

### Flow cytometric detection of MHCII, peptide/MHCII complexes, and associated machinery

To assess total surface MHCII expression in wild-type and knockout mice, cells were stained with a pan I-A/I-E anti-MHCII antibody. Human MHCII was detected using an anti-HLA-DR antibody. For detection of specific MHCII alleles in YAe presentation assays, cells were surface stained with anti-I-A^b^ and anti-I-E^d^ antibodies. Mouse invariant chain and H2-M expression were detected by staining intracellularly with anti-mouse CD74 and anti-H2-Mαβ2 antibodies, respectively. Two different fluorophore-conjugated versions of the anti-H2-Mαβ2 antibody were labeled in house, with the Pacific Blue and Alexa Fluor 647 labeling kits (P30013 or A20186, Thermo Fisher). Human invariant chain was detected using an anti-human CD74 antibody, and HLA-DM was detected using an anti-HLA-DM antibody^97^ provided by L. Denzin (Rutgers). CLIP-loaded MHCII molecules were detected on the cell surface by using an CLIP/I-A^b^ complex specific antibody^97^ provided by L. Denzin (Rutgers). To detect Eα^52-68^/I-A^b^ complexes, lung cells were surface stained with a biotinylated Y-Ae antibody followed by APC-Streptavidin (405207, Biolegend). Mouse H2-O expression was detected by staining intracellularly with an anti-mouse H2-Oβ antibody^97^ provided by L. Denzin (Rutgers).

### RNA isolation and quantitative PCR

Total RNA was extracted and purified using the Qiaqen RNeasy Plus Mini Kit (74134, Qiagen), and cDNA was then prepared using the SuperScript III First-Strand Synthesis System with random hexamers (18080051, Thermo Fisher). Quantitative PCR was performed using the Power SYBR Green PCR Master Mix system (4367659, Thermo Fisher) measured using a StepOnePlus Real Time PCR Machine (Applied Biosystems). Expression was quantified relative to the housekeeping gene *Hprt* using the ΔC_T_ method. The following forward (F) and reverse (R) primer pairs were used*. H2-Aa* F: CTGATTCTGGGGGTCCTCGC and R: CCTACGTGGTCGGCCTCAAT; *H2-Ab1* F: GAGCAAGATGTTGAGCGGCA and R: GCCTCGAGGTCCTTTCTGACTC; *H2-DMa* F: GGCGGTGCTCGAAGCA and R: TGTGCCGGAATGTGTGGTT; *H2-DMb1* F: CTATCCAGCGGATGTGACCAT and R: TGGGCTGAGCCGTCTTCT; *Hprt* F: TCAGTCAACGGGGGACATAA and R: GGGGCTGTACTGCTTAACCAG.

### Bone marrow chimeras

Donor bone marrow was isolated from C57Bl6 MHCII^-/-^ and CD45.1 B6 mice, and T cells were depleted magnetically using Thy1.2 Dynabeads (11443D, Thermo Fisher). 5×10^6^ bone marrow cells were then injected intravenously into lethally irradiated recipient C57Bl6 MHCII^-/-^ and CD45.1 B6 mice 6 hours after irradiation was completed (5.5 Gy x 2 doses, 3 hours apart). Mice were treated with Bactrim for 3 weeks following the transfer and were housed in autoclaved cages.

Full bone marrow chimera reconstitution was confirmed 6 weeks after transfer, by flow cytometry analysis of peripheral blood cells stained with anti-CD45.1 and anti-CD45.2 antibodies. MHCII expression on chimeric mouse lung cells was assessed 8 weeks post-transfer.

### DQ-ovalbumin assay

0.75-1.25×10^5^ AT2s, DCs, and B cells were incubated with either 0 or 10 μg/mL DQ-ovalbumin (D12053, Thermo Fisher) in 10% FBS in RPMI media at either 37°C or 4°C for 2 hours. Cells were then washed 3x with PBS and stained with Live/Dead Aqua; cells were maintained at 4°C during these steps. Fluorescence was then immediately assessed by flow cytometry.

### RNA-sequencing analysis

FASTQ files were obtained using the NCBI SRA toolkit from the GEO accession GSE115904 provided by Ma *et. al*^38^. Salmon quasi-alignment was then used to normalize and quantify gene expression to generate a transcripts per million (tpm) matrix^104^. This was performed using the “validateMappings” setting and the pre-computed partial mm10 index for Salmon provided on refgenie. The generated matrices were then imported into R v4.0.2 for conversion and concatenation of Ensembl transcript version IDs to gene symbols using the biomaRt package^105^.

### Cathepsin D and L activity assays

Cathepsin D (CtsD) activity was measured using the SensoLyte ® 520 Cathepsin D Assay Kit (AS-72170, Anaspec) per the manufacturer’s protocol, with lysates from 1×10^5^ cells plated in each well. Cathepsin L (CtsL) activity was measured using the SensoLyte ® Rh110 Cathepsin L Assay Kit (AS-72217, Anaspec) per the manufacturer’s protocol, with lysates from 4×10^5^ cells plated in each well.

For both assays, splenocytes were harvested as above, and AT2s were isolated via FACS, as above. For the CtsD activity assay, splenic B cells were isolated from splenocytes using the Dynabeads™ Mouse CD43 Untouched™ B Cells kit (11422D, Thermo Fisher), per the manufacturer’s protocol. For the CtsL activity assay, lung B cells were isolated via FACS, as above.

### Costimulatory molecule expression analysis

To generate bone marrow-derived dendritic cells (BMDCs), bone marrow cells were grown in R10 media supplemented with 20 ng/mL mouse recombinant granulocyte-macrophage colony-stimulating factor (“GM-CSF”; 10822-026, Shenandoah Biotechnology). Fresh media was added on days 3 and 6 after plating, and the floating fraction of cells was harvested on day 7-10.

Mice were infected intranasally with 60 FFU IAV PR8 diluted in 20 μL PBS under isoflurane anesthesia. Lungs were harvested from mice 5 days after infection or from naïve mice.

To evaluate the expression of costimulatory molecules, BMDCs, bulk splenocytes, and lung cells were surface stained with anti-CD80, anti-CD86, and anti-CD54 (ICAM1) antibodies.

### Detection of germline MHCII deletion

Consistent with prior reports^106, 107^, our Ab1^fl/fl^ mice experienced spontaneous germline disruption of the I-A^b^ locus at a rate of ∼5%, resulting in global loss of MHCII from all cells. Therefore, in addition to standard genotyping all SPC^ΔAb1^ and Ab1^fl/fl^ mice were screened at >5 weeks of age for germline deletion of MHCII, by flow cytometric measurement of MHCII on peripheral blood immune cells isolated by cheek bleed. Specifically, cells were stained for surface expression of CD45, CD19, and MHCII, and mice were excluded from further study if B cells (CD45^+^ CD19^+^) lacked MHCII expression. All mice included in experiments discussed here had appropriately intact MHCII expression by peripheral blood screening.

### Tamoxifen administration

SPC^ΔAb1^, Ab1^fl/fl^, and SPC^Cre^ mice >5 weeks of age were oral gavaged daily with 2 mg tamoxifen for 4 days, by delivering 100 μL of 20 mg/mL solution of tamoxifen (T5648, Sigma) in a 9:1 corn oil:ethanol mixture. Mice were used in experiments ≥5 days after the last dose of tamoxifen.

### Histological analysis

To assess MHCII expression in SPC^ΔAb1^, Ab1^fl/fl^, and MHCII^-/-^ mice by immunofluorescence, the lung vasculature was first perfused with 5 mL 4% paraformaldehyde in PBS (4% PFA) by injecting into the cardiac right ventricle. Lungs were then inflated via intratracheal instillation of 0.9 mL 4% PFA. The trachea was then tied with suture to keep the lungs inflated while they were excised en bloc and then placed in an additional 30 mL 4% PFA, then incubated for 24 hours at 4°C. Tissue was then processed, paraffin-embedded, sectioned, and stained with DAPI, and the following antibodies: anti-mouse pro-SPC (AB3786, EMD Millipore), anti-mouse MHCII (107601, Biolegend), and anti-mouse E-cadherin (ab76319, Abcam). Images were acquired using an Axio Observer 7 widefield microscope with Axiocam 702 monochrome CMOS camera and Zen blue acquisition software (Zeiss). Composite images were then generated in Fiji. To assess lung pathology at homeostasis in SPC^ΔAb1^ and Ab1^fl/fl^ mice, the lungs were inflated via intratracheal instillation of ∼1 mL 10% neutral buffered formalin (NBF), then placed in an additional ∼30 mL 10% NBF and incubated for 24 hours at RT. Tissue was then processed, paraffin-embedded, sectioned, and stained with hematoxylin and eosin, and the resulting lung sections were evaluated for signs of disease by a veterinary pathologist. Images were acquired using an Aperio AT2 Slide Scanner (Leica) and evaluated using Aperio ImageScope software (Leica).

### Homeostasis T cell phenotyping analyses

For determination of T cell phenotype as well as activation status, cells were surface stained for the following markers: CD44, CD62L, CD69, CD11a, PD1, and LAG3. Cells were also stained intranuclearly for FoxP3 and Ki67.

### Lung organoids

Alveolar organoids were cultured as described previously^29, 31^, with the following modifications. Briefly, 5×10^3^ AT2s sorted from either SPC^ΔAb1^ and Ab1^fl/fl^ mice were cocultured with 5×10^4^ Pdgfrα^+^ lung fibroblasts sorted from WT C57Bl/6 mice. Cells were suspended in 90μL of a 1:1 mixture of Matrigel:modified SAGM media (CC-3118, Lonza). SAGM media is modified by adding 0.1 μg/mL cholera toxin (C8052, Sigma) and the following SAGM BulletKit components: 10 μg/mL insulin, 5 μg/mL transferrin, 25 ng/mL EGF, 30 μg/mL bovine pituitary extract, 0.01μM retinoic acid, and 5% FBS. The 1:1 mixture containing cells was placed in a cell culture insert (353095, Corning); after solidification of the matrix at 37°C, the insert was placed inside a well of 24-well plate containing modified SAGM media. For the first two days of culture, 10 μM ROCK inhibitor Y27632 (Y0503, Sigma) was added to the media. The media was changed every 48 hours, and after 21 days organoids were imaged using an EVOS FL Auto. Organoid numbers were quantified in Fiji using the Cell Counter plugin.

### Virus infections for weight loss and mortality analysis

For influenza infection weight loss monitoring, mice were infected intranasally with 3 FFU IAV PR8 diluted in 20 μL PBS under isoflurane anesthesia. For influenza infection mortality experiments, mice were infected intranasally with 15 FFU IAV PR8 diluted in 20 μL PBS under isoflurane anesthesia. For both Sendai virus infection weight loss monitoring and mortality analyses, mice were infected intranasally with 3.5-7×10^4^ TCID_50_ SeV 52 diluted in 35 μL PBS under ketamine (70 mg/kg) + xylazine (5 mg/kg) anesthesia. For all weight loss experiments, mice were weighed on the days indicated. For all mortality studies, mice were monitored daily for death or moribund state as an endpoint.

### Lung virus titering

For influenza virus titering, mice were infected intranasally with 3 FFU IAV PR8 diluted in 35 μL PBS under ketamine (70 mg/kg) + xylazine (5 mg/kg) anesthesia. For Sendai virus titering, mice were infected intranasally with 7×10^4^ TCID_50_ SeV 52 diluted in 35 μL PBS under ketamine (70 mg/kg) + xylazine (5 mg/kg) anesthesia. Mice were sacrificed at the timepoints indicated for virus titers analysis.

For influenza virus titers determination, MDCK cells were infected with serial 10-fold dilutions of lung homogenates in MEM containing 50μg/mL gentamicin, 5mM HEPES, and 1μg/mLTPCK-treated trypsin (LS003750, Worthington Biochemical) in quadruplicate per lung. After 4 days of incubation at 37°C, the presence of virus at each dilution was determined by visual cytopathic effect, and lung virus titers were determined using the Reed and Muench Calculation.

For Sendai virus titers determination, LLC-MK2 cells were infected with serial 10-fold dilutions of lung homogenates in DMEM containing 50μg/mL gentamicin, 0.35% BSA, 0.12% NaHCO3, and 2 μg/mL TPCK-treated trypsin in triplicate per lung. After 3 days of incubation at 37°C, the presence of virus was determined by assessing the capacity of 25μL of supernatant at each dilution to hemagglutinate 0.25% chicken red blood cells (cRBCs) in a 100μL final volume after a 30” incubation at room temperature. Lung virus titers were then determined using the Reed and Muench Calculation.

### Influenza LAG3^+^ T cell expansion analysis

Mice were infected intranasally with 3 FFU IAV PR8 diluted in 20 μL PBS under isoflurane anesthesia. Mice were sacrificed 9 days after infection and lungs were stained *ex vivo* for T cell surface markers and LAG3.

### Influenza lung PD-L1 and CD44, CD62L, and PD1 expression analysis

Mice were infected intranasally with 3 FFU IAV PR8 diluted in 20 μL PBS under isoflurane anesthesia. For PD-L1 analysis, mice were sacrificed 9 days after infection and lungs were dissociated for AT2s as above, then stained *ex vivo* for AT2 cell markers and PD-L1. For PD1, CD44, CD62L analysis, mice were sacrificed 9 days after infection and lungs were dissociated for T cells as above, then stained *ex vivo* for T cell surface markers, PD1, CD44, and CD62L.

### Hybridoma assay

NFAT-*lacZ*-inducible T cell hybridomas recognizing MHCII-restricted influenza-derived epitopes have been described previously^54, 63^. Hybridoma recognition of cognate peptide/MHCII complexes results in β-galactosidase production, which was detected using a fluorometric β-galactosidase substrate 4-methyl-umbelliferyl-β-D-galactopyranoside (“MUG”; M1633, Sigma).

1×10^4^ APCs were treated with media only, 20 μg/mL peptide, or 1×10^6^ FFU influenza virus in R10 media for 45” at 37°C, 6% CO_2_ in a 384 well plate; for AT2 conditions, wells were pre-coated with a thin layer of 9:1 R10:Matrigel (356231, Corning) mixture. After 45”, 2×10^4^ hybridomas were directly added, and cells were then cocultured for 14-18 hours at 37°C, 6% CO_2_. MUG substrate solution (33 μg/mL MUG in PBS containing 38.5 μM 2-mercaptoethanol, 9 mM MgCl2, and 1.25% NP40) was then added to the co-culture at a ratio of 1:5 MUG solution:co-culture, then incubated for 3 hours at 37°C, 6% CO_2_. Fluorescence (excitation: 365 nm, emission: 445nm) was detected on Tecan Infinite M200 Pro Plate Reader.

### IFNγ ELISpot assay

For isolation of influenza-infected APCs, WT and MHCII^-/-^ mice were infected intranasally with 1.2×10^6^ FFU IAV PR8 diluted in 35 μL PBS under ketamine (70 mg/kg) + xylazine (5 mg/kg) anesthesia. Lungs were harvested from mice 4 days after infection or from naïve mice. APCs were isolated via cell sorting, as above.

For isolation of influenza-experienced effector T cells, WT mice were infected intranasally with 3 FFU IAV PR8 diluted in 35 μL PBS under ketamine (70 mg/kg) + xylazine (5 mg/kg) anesthesia. Spleens were harvested from mice 10 days after infection. CD4^+^ and CD8^+^ T cells were isolated from bulk splenocytes using Dynabeads Untouched Kits for mouse CD4 cells and CD8 cells, respectively, per the manufacturer’s protocol (11415D and 11417D, Thermo Fisher).

96-well MultiScreen_HTS_ IP 0.45 μm filter plates (MSIPS4W10, Millipore-Sigma) were coated with mouse IFNγ capture antibody (551881, BD) and incubated at 4°C for 20 hours prior to the assay. Plates were blocked with R10 media for >1 hour at 37°C, after which 5×10^4^ APCs and 1×10^5^ T cells were co-cultured for 14-18 hours in the presence of 2μg/mL anti-CD28 (40-0281-M001, Tonbo Biosciences). T cell IFNγ production was detected via biotinylated IFNγ detection antibody (551881, BD), followed by HRP-Streptavidin (557630, BD) and chromogenic 3-Amino-9-ethylcarbazole (AEC) substrate (551951). IFNγ spots were imaged and counted using a CTL ImmunoSpot S6 Universal Analyzer.

### Detection of influenza-infected cells

For quantification of influenza virus infection in hybridoma and ELISpot assays, at assay endpoint cells were stained for surface hemagglutinin (HA) protein expression or intracellular nucleoprotein (NP) protein expression. Two different fluorophore-conjugated versions of the anti-HA antibody were labeled in house, with the Alexa Fluor 488 and Alexa Fluor 647 labeling kits (A20181 or A20186, Thermo Fisher).

### IFNγ treatment of mice

Mice were injected intravenously under isoflurane anesthesia once daily with 1×10^5^ U recombinant mouse IFNγ (575306, Biolegend) in 150 μL PBS, or PBS only, for 3 days in a row.

### Assay schematics

Assay diagrams in Figs 5-6 were adapted from images created with BioRender.com.

### Statistical analysis

All statistical analyses were performed using Prism 8 software (GraphPad). The statistical tests used are indicated in the figure legend corresponding to each specific experiment. Unpaired two-tailed Student’s *t*-test was used to compare means between two groups if they had similar variances by F-test, and if they passed all four of the following normality of residuals tests: Anderson-Darling, D’Agostino-Pearson omnibus, Shapiro-Wilk, and Kolmogorov-Smirnov. If the samples failed at least one normality test then two-tailed Mann-Whitney test was used. For weight loss analysis, a mixed model, which uses a compound symmetry covariance matrix and is fit using Restricted Maximum Likelihood (REML), with Geisser-Greenhouse correction was used, followed by post-hoc multiple comparisons with Sidak’s correction; this was used instead of repeated-measures ANOVA, as it can handle missing values. For mortality analysis, the Log-rank (Mantel-cox) test was used to compare survival curves. Two-way ANOVA followed by post-hoc multiple comparisons with Tukey’s or Sidak’s correction was used to compare means between groups with 2 contributing independent variables; Tukey’s correction was used when all possible means were compared, and Sidak’s correction was used when means were compared across one factor only. Three-way ANOVA followed by post-hoc multiple comparisons with Sidak’s correction was used to compare means between groups with 3 contributing independent variables.

### Data availability

All original data generated or analyzed in this study are available from the corresponding author upon reasonable request.

## Acknowledgements

We thank the Children’s Hospital of Philadelphia (CHOP) Flow Cytometry Core, in particular F. Tuluc, for their assistance with the flow cytometry and cell sorting experiments. We thank E. Radaelli and the Penn Vet Pathology Core for their help with our tissue histology studies, and we thank A. Stout and the Penn CBD Microscopy Core for their help with imaging. We thank the CHOP Abramson Research Center Animal Facility staff for their assistance with mouse colony maintenance. We thank C. Lopez and D. Fisher (University of Pennsylvania) for their assistance with the Sendai virus work and L. Denzin (Rutgers) for providing reagents and helpful discussion. We thank J. Ma and colleagues (University of Melbourne) for depositing their RNA-sequencing FASTQ files on GEO. The work was supported by National Institutes of Health grants 5R01AI113286 (L.C.E), T32AI007324 (S.A.T), F30HL145907 (S.A.T), and the CHOP Division of Protective Immunity and Immunopathology Pilot Grant (S.A.T).

## Author Contributions

S.A.T. and L.C.E. conceptualized the project. S.A.T. designed and performed all of the experiments, except for the RNA-seq analysis (Fig S5a), which was conducted by J.H.L., and the growth of organoids (Fig S8), which was conducted by A.J.P. L.C.E. supervised and provided guidance for all studies. C.B. provided critical assistance with the *in vivo* mouse studies. A.J.P. and G.S.W. contributed essential guidance for the study and isolation of AT2s and provided the SPC-Cre-ERT2 mice. J.K, M.C.B, and E.E.M. established the pipeline through which human AT2s were acquired (Fig S2). S.A.T. analyzed the data, made the figures, and wrote the original draft. L.C.E edited the manuscript and figures. All authors reviewed the manuscript.

---

**Supplementary Figure 1:**
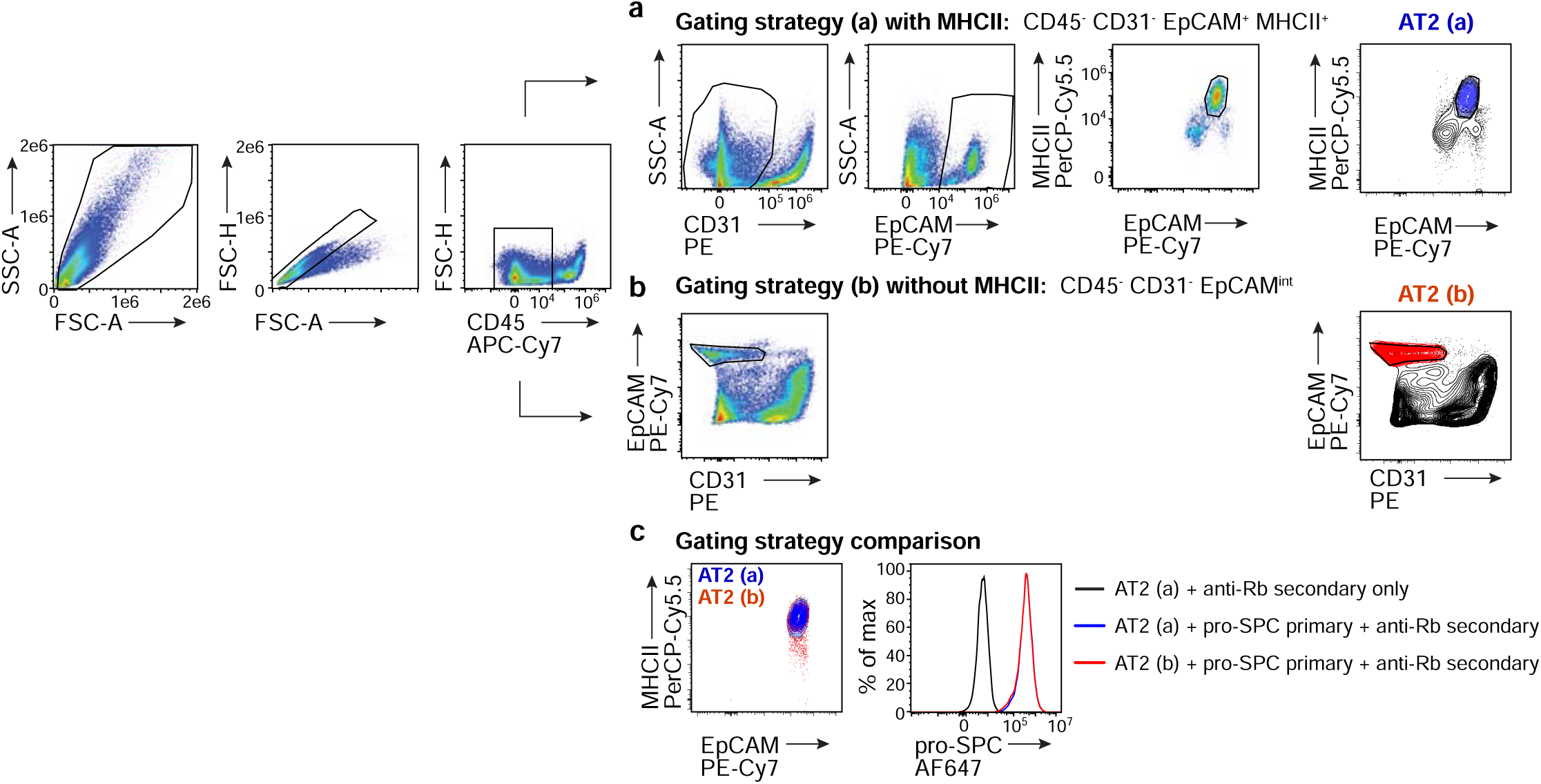
Murine AT2s gating strategies. **a,b,** General gating strategy for cell sorting of murine AT2s using MHCII as a positive marker (**a**) or without using MHCII as a positive marker (**b**). The right column contour plots are the same as the final pseudocolor plots in the corresponding gating strategies, and they are shown to highlight the final AT2 population defined from each strategy. **c,** Comparison of the two populations of AT2s identified by the two gating strategies in terms of MHCII expression (contour plot, left) or pro-SPC expression (histogram, right).

**Supplementary Figure 2:**
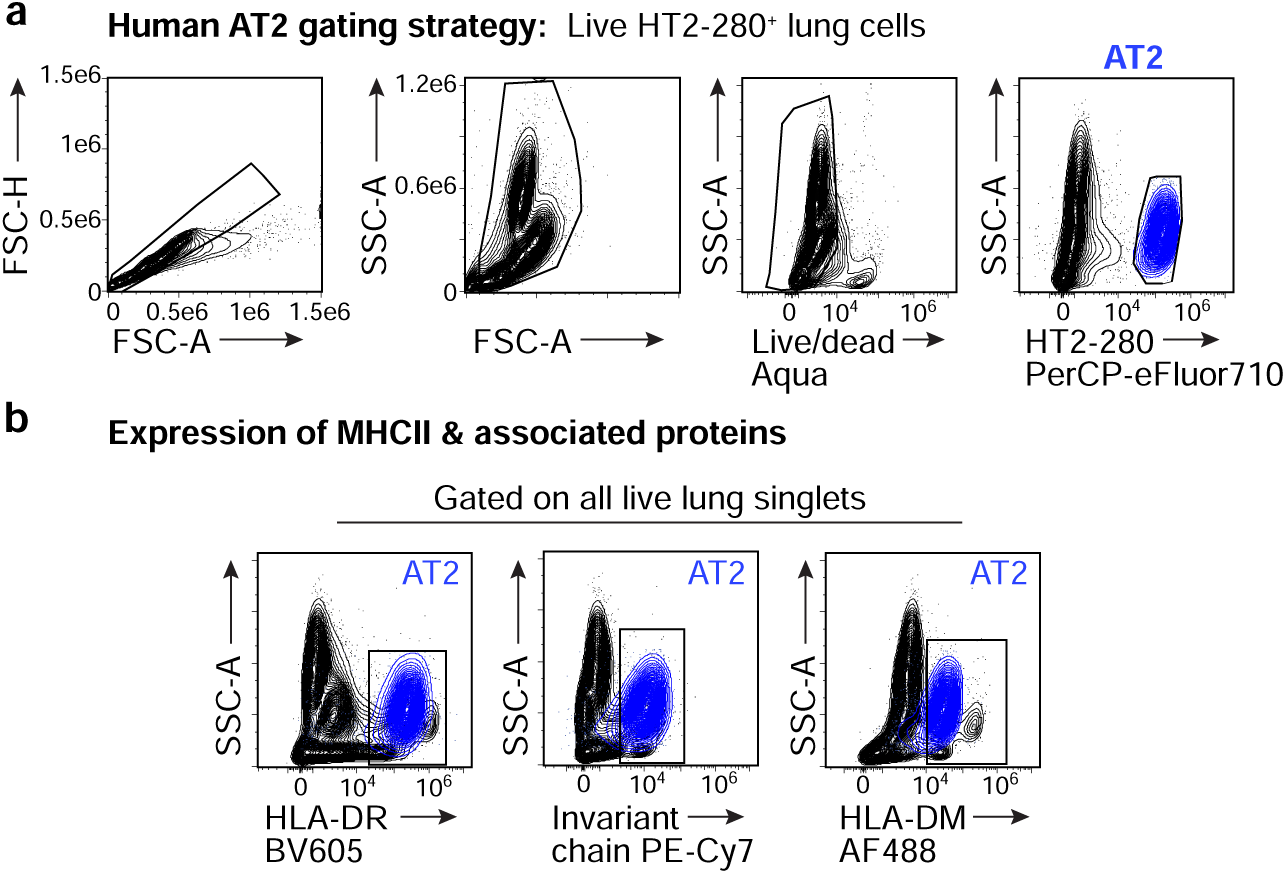
Human AT2s express MHCII and associated classical MHCII presentation mediators. **a,** Gating strategy for human AT2s. The right-most contour plot highlights the final AT2 population defined by HT2-280^+^ staining. **b,** HLA-DR (left), invariant chain (middle), and HLA-DM (right), protein expression by healthy human distal lung cells (black), quantified by *ex vivo* flow cytometry analysis. The AT2 population, as defined in (**a**), is highlighted in blue. Gates were drawn based on fluorescence minus one (FMO) controls. All plots represent n=1 human donor.

**Supplementary Figure 3:**
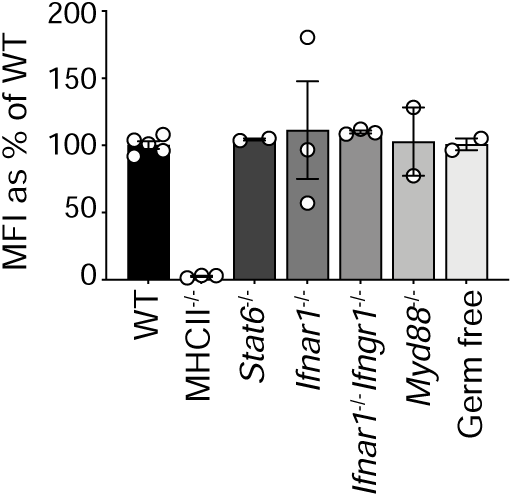
AT2s constitutively express MHCII independent of a variety of inflammatory mediators. MHCII protein expression by AT2s, from naïve B6 WT, MHCII^-/-^ *Stat6*^-/-^, *Ifnar1*^-/-^, *Ifnar1*^-/-^ *Ifngr1*^-/-^, *Myd88*^-/-^, and germ-free mice, measured by *ex vivo* flow cytometry analysis. For each strain, MHCII expression is displayed as a percentage reflecting the average median fluorescence intensity (MFI) of MHCII on AT2s relative to the MHCII MFI on AT2s in WT mice; symbols indicate n=2-5 mice per strain, pooled from 1-2 independent experiments. Data are shown as mean plus standard error of the mean (SEM).

**Supplementary Figure 4:**
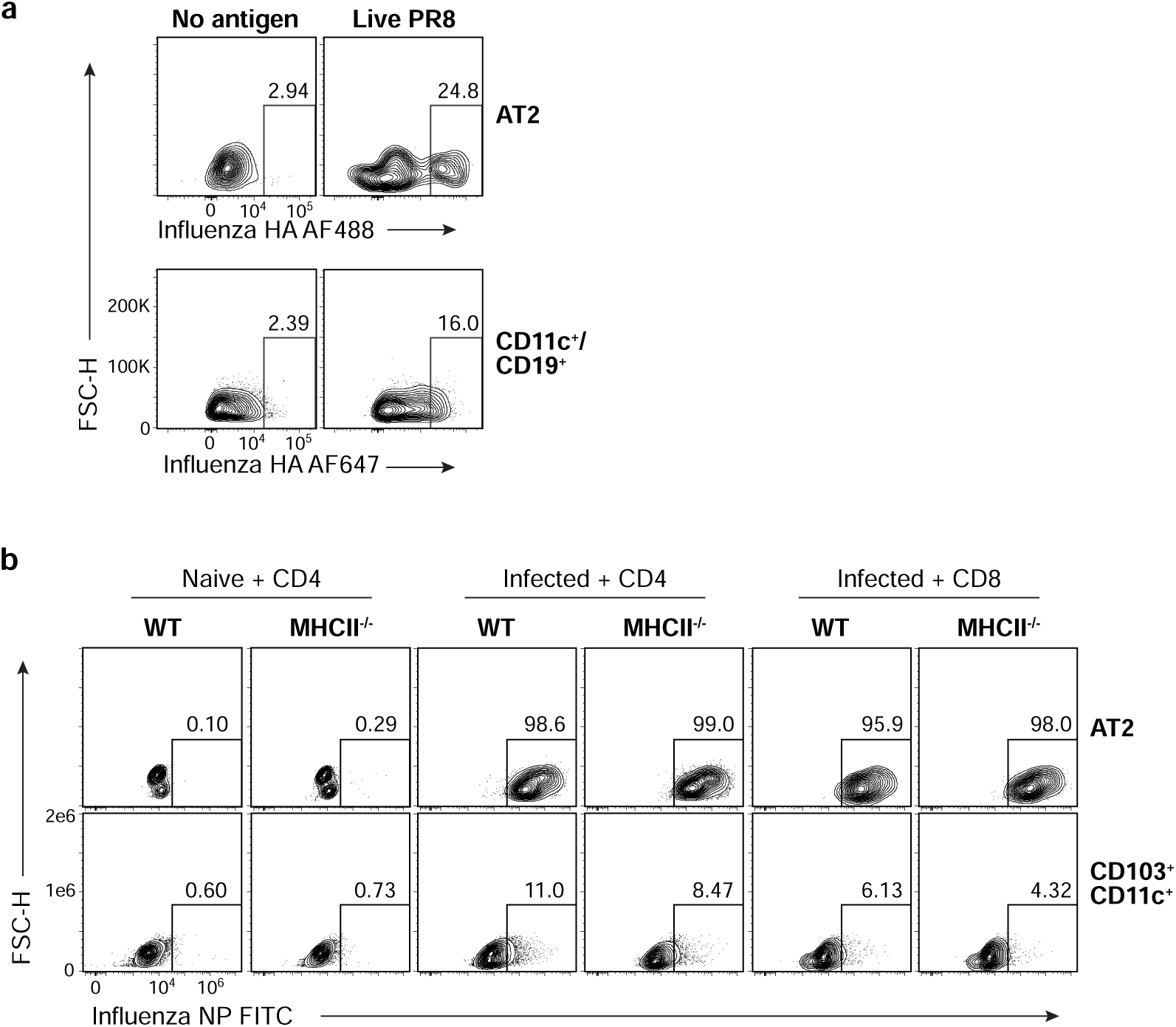
AT2s are infected robustly with influenza *in vitro* and *in vivo*. **a,** Surface influenza hemagglutinin (HA) expression by B6 AT2s (top) and a mixed population of CD11c^+^ and CD19^+^ lung cells (bottom) sorted from naïve mice then incubated with no antigen (left) or live virus (right) *in vitro* for 14h. These plots reflect the antigen presenting cells used in the hybridoma antigen presentation assay illustrated in Figure 5b. The frequency of HA^+^ cells is shown above the gates, which were drawn based on the no antigen conditions. Two different fluorophore conjugated versions of the same anti-HA antibody were used because the AT2 and CD11c^+^/CD19^+^ populations were already labeled with different fluorophores (APC and FITC, respectively) from the cell sorting process. **b,** Intracellular influenza nucleoprotein (NP) expression by B6 AT2s (top) and CD103^+^CD11c^+^ lung cells (bottom) sorted from naïve or flu-infected WT or MHCII^-/-^ B6 mouse lungs 4 dpi and incubated with CD4^+^ and CD8^+^ T cells *in vitro* for 14h (as indicated). These plots reflect the antigen presenting cells used in the ELISpot presentation assay illustrated in Figure 5e. The frequency of NP^+^ cells is shown above the gates, which were drawn based on the naïve conditions.

**Supplementary Figure 5:**
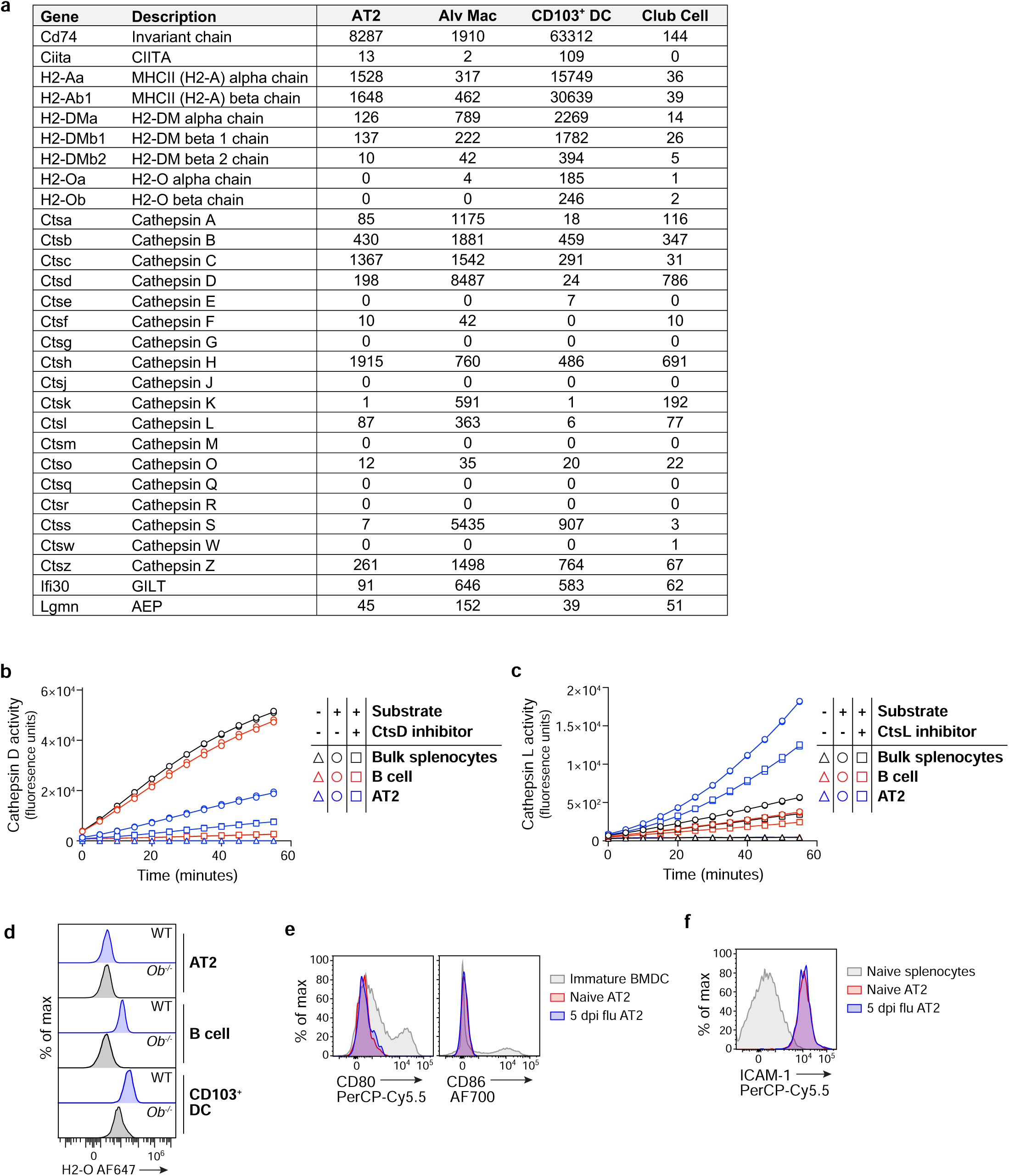
AT2s express conventional mediators of MHCII antigen processing and presentation as well as the noncanonical costimulatory molecule ICAM-1, but do not express CD80, CD86, or the H2-M antagonist H2-O. **a,** Expression of MHCII-associated genes and proteases in naïve B6 WT AT2s, alveolar macrophages, CD103^+^ DCs, and club cells, evaluated by RNA-sequencing analysis of data from Ma *et. al.*^38^ Expression values are reported as transcripts per million (TPM) and represent the average of n=4 biological replicates for each cell type. **b,** Cathepsin D (CtsD) activity of AT2s, bulk splenocytes, and splenic B cells from naïve B6 mice measured by cleavage of a fluorogenic CtsD substrate in the presence or absence of the CtsD inhibitor pepstatin A (10 nM); symbols shown are 2 technical replicates from 1 biological replicate and are representative of n=2-3 biological replicates per cell type. **c,** Cathepsin L (CtsL) activity of AT2s, bulk splenocytes, and lung B cells isolated from naïve B6 mice, measured by cleavage of a fluorogenic CtsL substrate in the presence or absence of a CtsL inhibitor (100 nM); symbols shown are 2 technical replicates from 1 biological replicate, and are representative of n=1-4 biological replicates per cell type. **d,** Intracellular H2-O protein expression by AT2s, CD103^+^ DCs, and B cells, from B6 WT or H2-Oβ deficient (*Oβ*^-/-^) mouse lungs measured *ex vivo* by flow cytometry; histograms represent n=2-3 mice per strain from 1 experiment. **e,** Surface expression of CD80 (left) and CD86 (right) by AT2s from naïve mice or mice 5 days post influenza virus infection, and B6 WT BMDCs cultured *in vitro*. **f,** Surface expression of ICAM-1 by AT2s from naïve mice or mice 5 days post influenza virus infection, and B6 WT splenocytes from naïve mice. Histograms represent n=2 mice (naïve) or n=6 mice (flu infected) in total across 2 experiments.

**Supplementary Figure 6:**
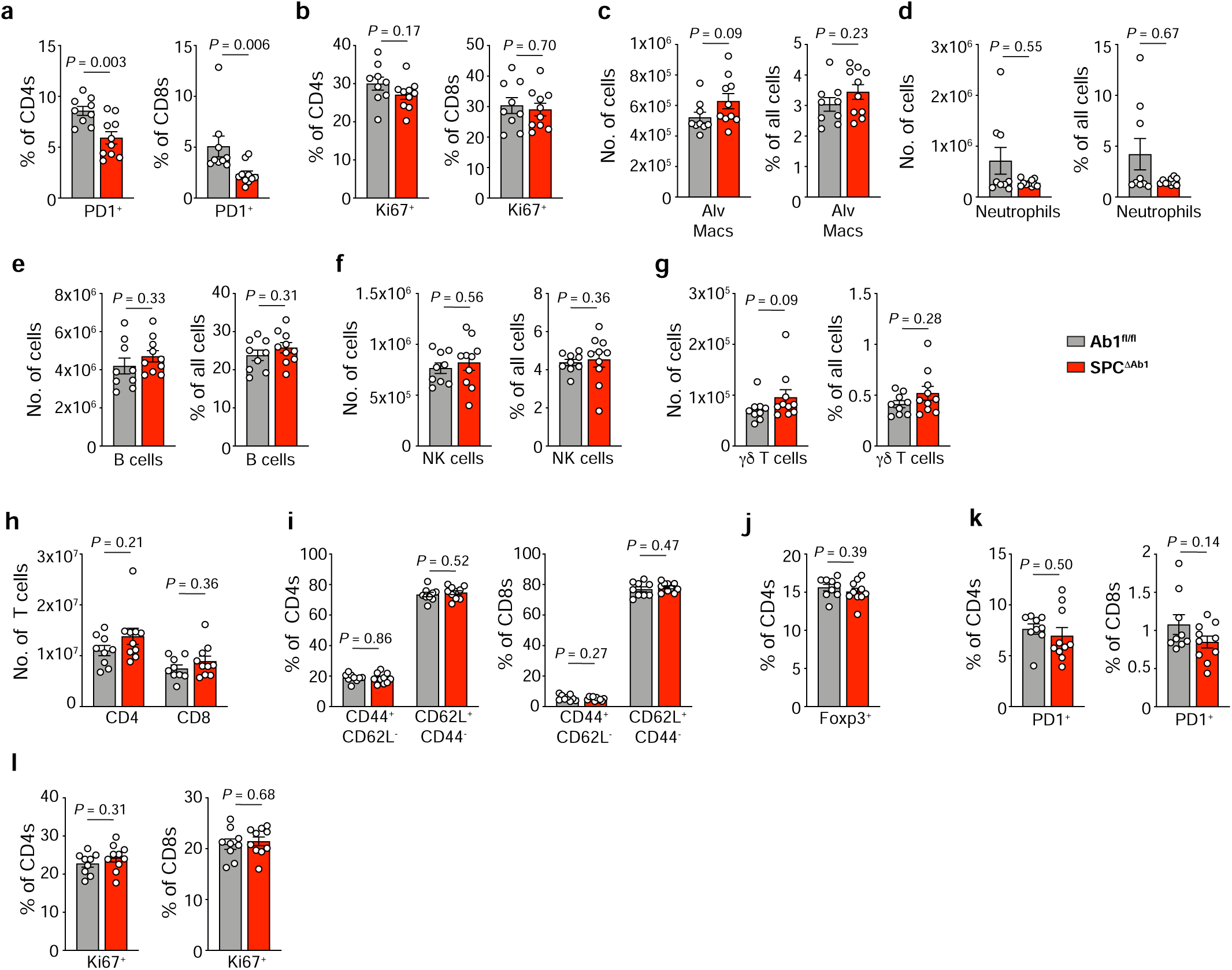
Mice lacking AT2 MHCII demonstrate normal immune homeostasis in the lung and spleen. *Ex vivo* flow cytometry analysis of naïve 12 week old Ab1^fl/fl^ and SPC^ΔAb1^ mice lungs and spleens. **a-b,** Frequency of lung CD4^+^ and CD8^+^ T cells expressing PD1 (**a**), and Ki67 (**b) c-g,** Absolute numbers and frequencies of lung alveolar macrophages (**c**), neutrophils (**d**), B cells (**e**), NK cells (**f**), γδT cells (**g**). **h-l**, Absolute number of splenic CD4^+^ and CD8^+^ T cells (**h**), and frequency of splenic CD4^+^ and CD8^+^ T cells expressing CD44, CD62L (**i**), Foxp3 (**j**), PD1 (**k**), and Ki67 (**l**). **a-l**, Data shown are mean plus SEM of n=9-10 mice per strain from 1 experiment, representative of 2 independent experiments, analyzed by unpaired two-tailed Student’s *t*-test (**a** [CD4s], **b, c** [%], **e**, **f** [#], **i-l**) or Mann-Whitney test (**a** [CD8s], **c** [#], **d**, **f** [%]**, g, h**). Full statistical test results are in Extended Data Table 1.

**Supplementary Figure 7:**
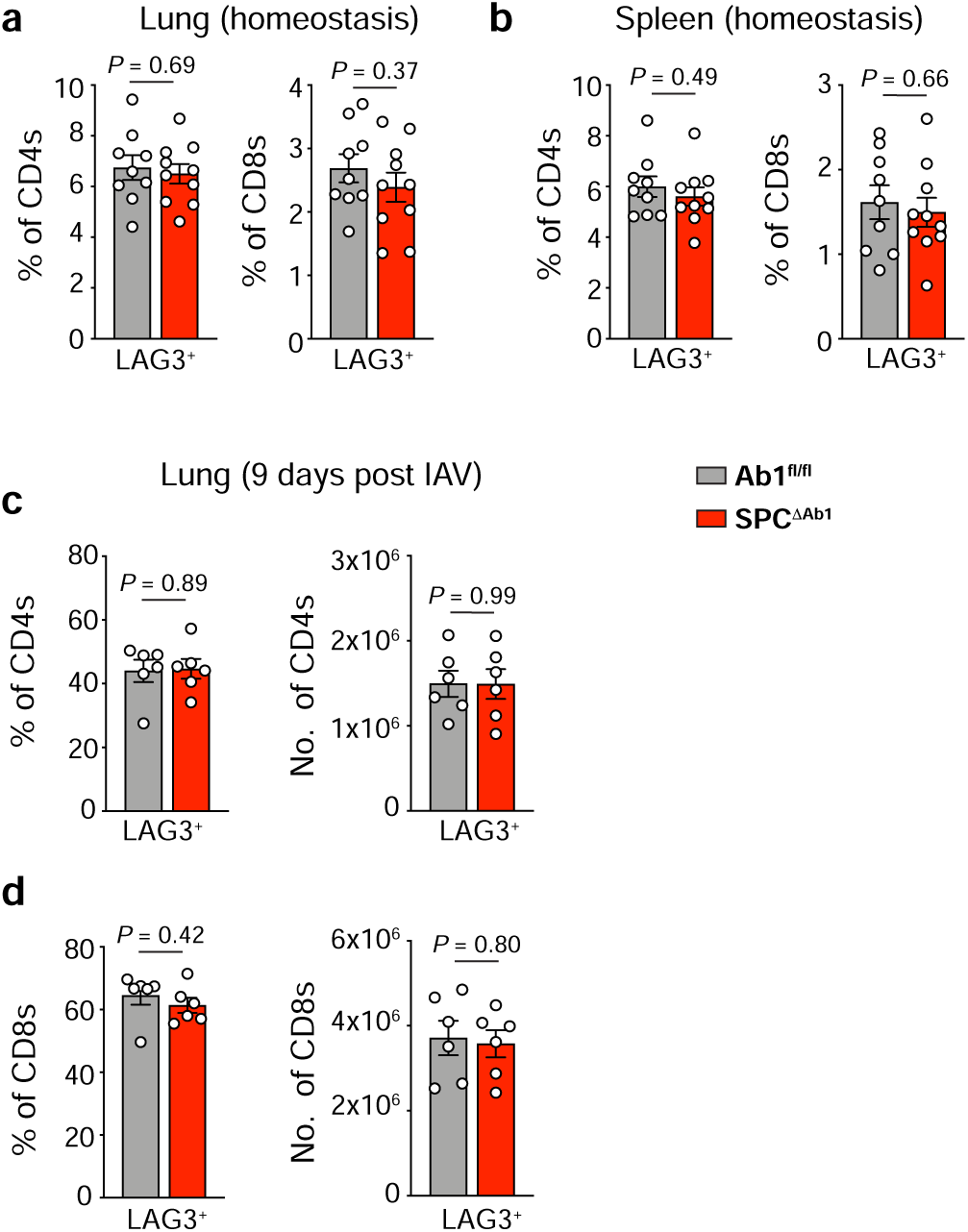
Loss of AT2 MHCII does not alter frequencies of LAG3+ T cells at homeostasis or after influenza virus infection. *Ex vivo* flow cytometry analysis of LAG3 expression by CD4^+^ and CD8^+^ T cells in Ab1^fl/fl^ and SPC^ΔAb1^ mice. **a,b,** Frequency of lung (**a**) and spleen (**b**) LAG3^+^ CD4^+^ and CD8^+^ T cells in naïve 12 week old mice; n=9-10 mice per strain from 1 experiment, representative of 2 independent experiments. **c,d,** Frequency and number of lung LAG3^+^ CD4^+^ (**c**) and CD8^+^ (**d**) T cells in 8-10 week old mice 9 days post IAV infection; n=6 mice per strain from 1 experiment. **a-d**, Data shown are mean plus SEM, and were analyzed by unpaired two-tailed Student’s *t*-test. Full statistical test results are in Extended Data Table 2.

**Supplementary Figure 8:**
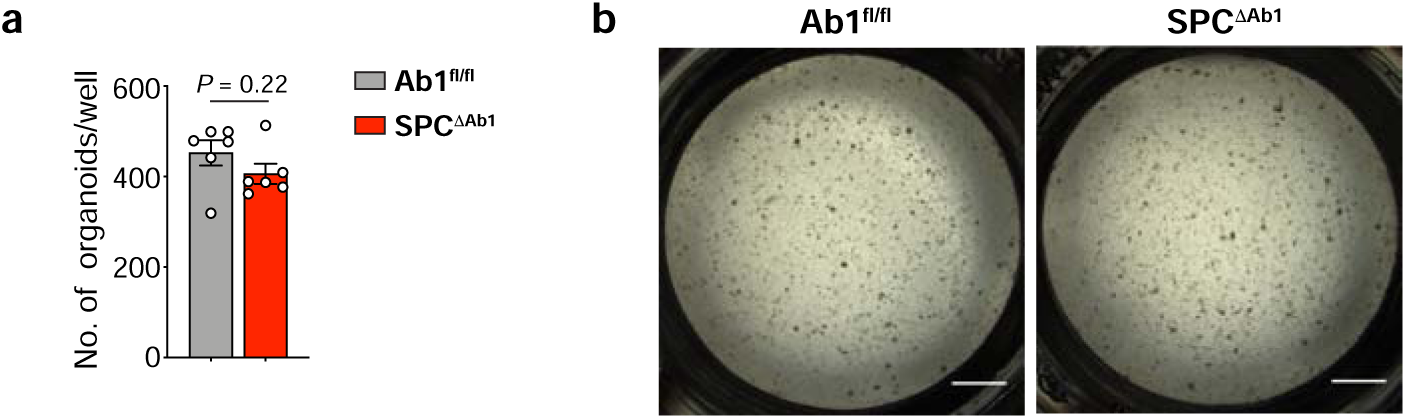
Loss of MHCII from AT2s does not inhibit their regenerative capacity. **a, b,** Numbers (**a**) and representative images (**b**) of lung organoid cultures formed by AT2s sorted from either or Ab1^fl/fl^ and SPC^ΔAb1^ mice. Data represent n=6 organoid cultures per strain, grown from AT2s originally sorted from n=1 mouse per strain. **a,** Bars represent mean plus SEM, and the data were analyzed by unpaired two-tailed Student’s *t*-test (t=1.294, df=10). **b,** Scale bar depicts 1000 μm.

**Supplementary Figure 9:**
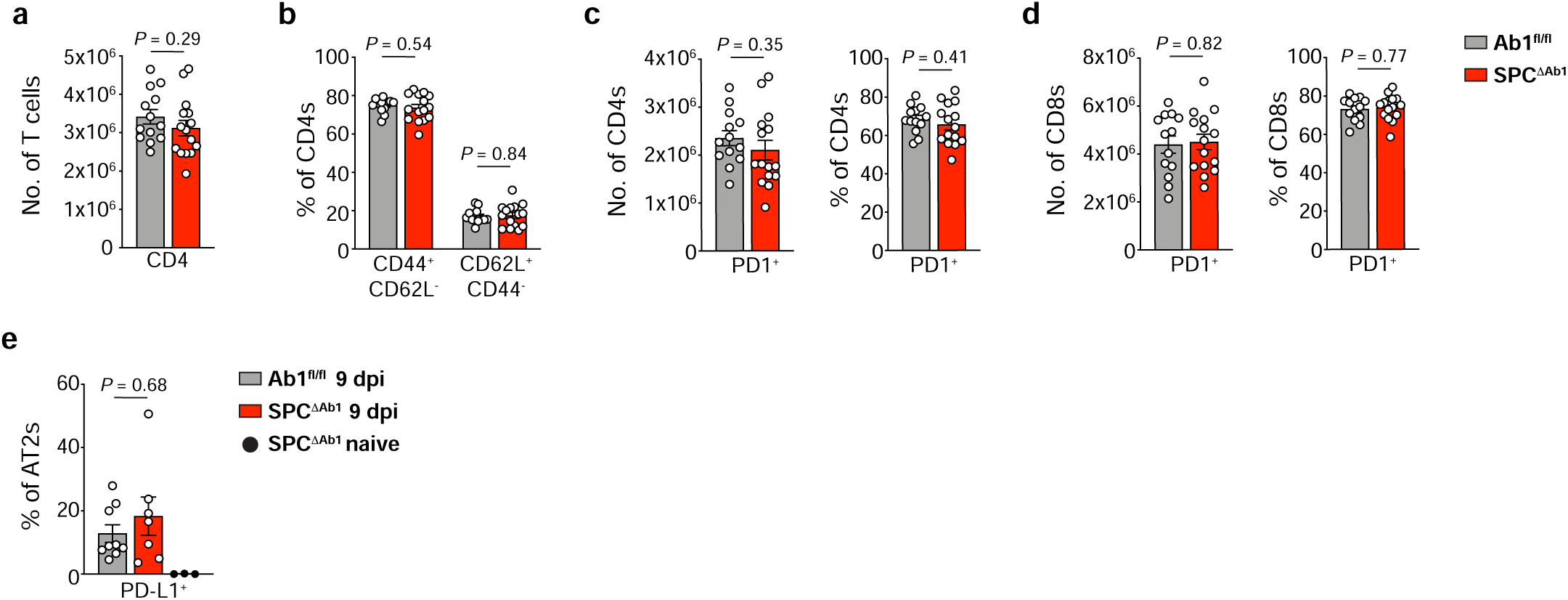
Loss of AT2 MHCII does not alter lung bulk or CD44/CD62L-expressing CD4+ T cells, PD1-expressing T cells, or PD-L1+ AT2s after influenza virus infection. *Ex vivo* flow cytometry analysis of PD-L1 expression by AT2s, and CD44, CD62L, and PD1 expression by lung CD4^+^ and CD8^+^ T cells. **a-d,** Number of bulk lung CD4+ T cells (**a**), frequency of CD4^+^ T cells expressing CD44, CD62L (**b**), and number and frequency of lung PD1^+^ CD4^+^ (**c**) and CD8^+^ (**d**) T cells in Ab1^fl/fl^ and SPC^ΔAb1^ mice 9 days post IAV infection; n=13-15 mice per strain, combined from 2 independent experiments. **e,** Frequency of PD-L1^+^ AT2s in Ab1^fl/fl^ and SPC^ΔAb1^ mice 9 days post IAV infection and naïve SPC^ΔAb1^ mice; n=7-9 mice per strain post-IAV and n=3 naïve, combined from 2 independent experiments. **a-e**, Data shown are mean plus SEM, and were analyzed by unpaired two-tailed Student’s *t*-test (**a, b**[CD44^-^CD62L^+^]**, c, d**), Welch’s *t*-test (**b** [CD44^+^CD62L^-^]), and Mann-Whitney test (**e**). Full statistical test results are in Extended Data Table 5.

**Supplementary Table 1:** List of flow cytometry antibodies used.

| **Flow cytometry antibodies used** |  |  |
| --- | --- | --- |
| **Target** | **Clone or Catalog #** | **Source** |
| mouse CD45 | 30-F11 | Biolegend |
| mouse CD31 | 390 | Biolegend |
| mouse EpCAM (CD326) | G8.8 | Biolegend |
| mouse I-A/I-E (MHCII) | M5/114.15.2 | Biolegend |
| mouse proSP-C | AB3786 | Sigma Aldrich |
| human HT2-280 | TB-27AHT2-280 | Terrace Biotech |
| mouse IgM | II/41 | Thermo Fisher |
| mouse CD11c | N418 | Biolegend |
| mouse CD103 | 2E7 | Biolegend |
| mouse CD11b | M1/70 | BD |
| mouse CD64 | X54-5/7.1 | Biolegend |
| mouse CD3 | 145-2C11 or 17A2 | BD or Biolegend |
| mouse CD8 | 53-6.7 | BD |
| mouse CD19 | 6D5 | Biolegend |
| mouse B220 | RA3-6B2 | Biolegend |
| mouse CD4 | RM4-5 or GK1.5 | Biolegend |
| mouse NK1.1 | PK136 | Thermo Fisher |
| mouse TCRδ | GL3 | Biolegend |
| mouse Ly6G | 1A8 | BD |
| mouse Podoplanin | 8.1.1 | Thermo Fisher or Biolegend |
| mouse Sca-1 | D7 | Thermo Fisher |
| mouse CD34 | MEC14.7 | Biolegend |
| mouse Pdgfrα (CD140a) | APA5 | Thermo Fisher |
| human HLA-DR | G46-6 | BD |
| mouse I-A<sup>b</sup> | AF-120.1 | Thermo Fisher |
| mouse I-E<sup>d</sup> | 14-4-4S | Biolegend |
| mouse invariant chain (CD74) | In-1 | BD |
| mouse H2-Mα β2 | 2E5A | BD |
| mouse H2-Oβ | Mags.Ob1 | L. Denzin |
| human invariant chain (CD74) | Pin.1 | Biolegend |
| human HLA-DM | Map.DM1 | L. Denzin |
| mouse CLIP/I-A<sup>b</sup> | 15G4 | L. Denzin |
| mouse Eα<sup>52-68</sup>/I-A<sup>b</sup> | YAe | Thermo Fisher |
| mouse CD80 | 16-10A1 | BD |
| mouse CD86 | GL1 | BD |
| mouse ICAM-1 (CD54) | YN1/1.7.4 | Biolegend |
| mouse CD45.1 | A20 | Biolegend |
| mouse CD45.2 | 104 | Biolegend |
| influenza hemagglutinin (HA) | IC5-4F8 | BEI |
| influenza nucleoprotein (NP) | D67J | Thermo Fisher |
| mouse CD44 | IM7 | Biolegend |
| mouse CD62L | MEL-14 | Biolegend |
| mouse CD69 | H1.2F3 | Biolegend |
| mouse CD11a | M17/4 | Biolegend |
| mouse PD1 | RMP1-30 | Biolegend |
| mouse LAG3 | C9B7W | Biolegend |
| mouse Foxp3 | FJK-16s | Thermo Fisher |
| mouse Ki67 | 16A8 | Biolegend |
| mouse PD-L1 | MIH5 | BD |

**Extended Data Table 1:**
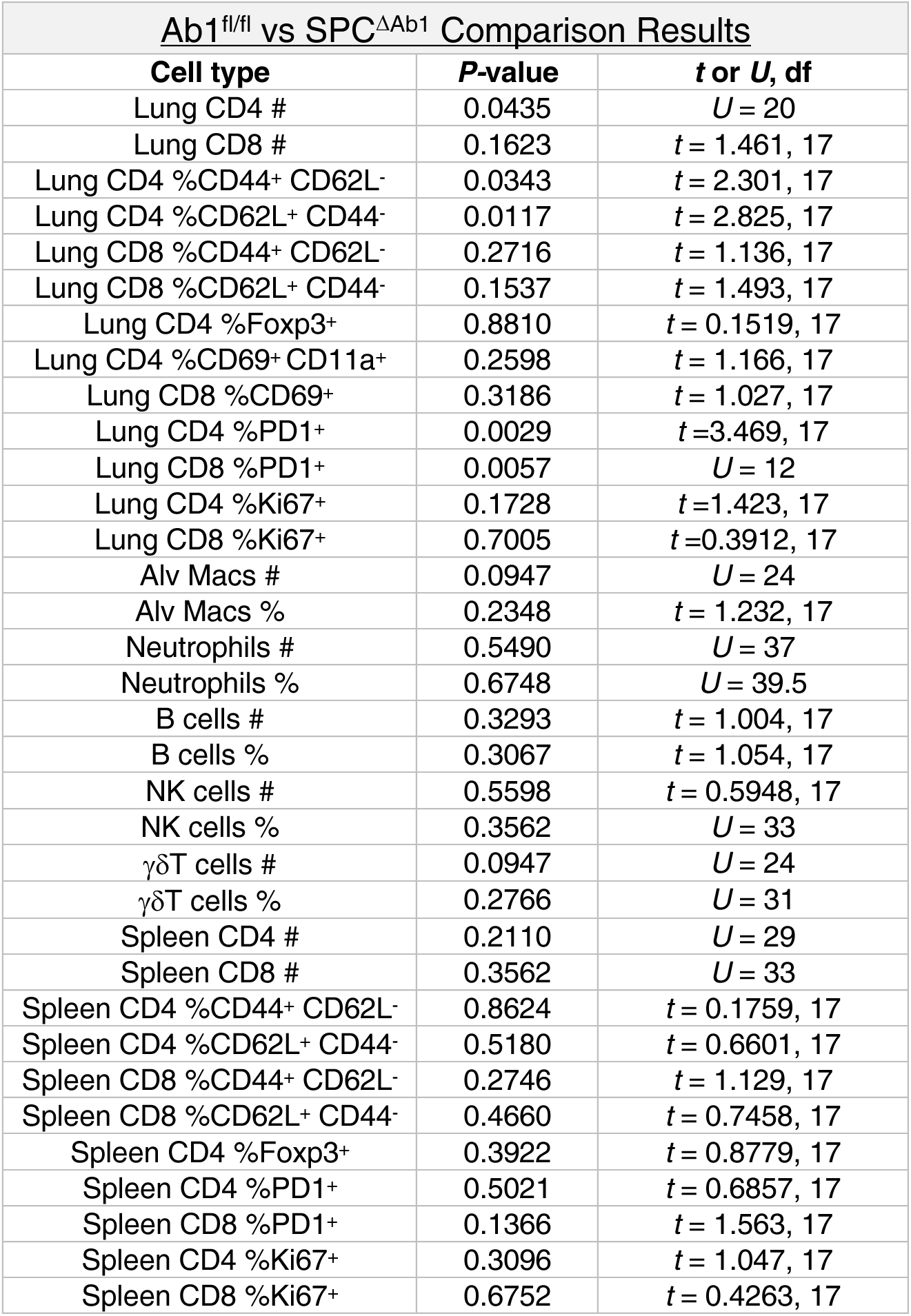
Statistical test results for comparisons of lung and spleen immune cells at homeostasis in Ab1^fl/fl^ and SPC^ΔAb1^ mice. Results of unpaired two-tailed Student’s *t*-tests and Mann-Whitney tests comparing immune cell populations between Ab1^fl/fl^ and SPC^ΔAb1^ mice. *P* values and associated *U* or *t* statistics with associated degrees of freedom (df) for each comparison are reported. Statistical results in this table are for the data depicted in Figures 3f-i and S6.

**Extended Data Table 2:** Statistical test results for comparisons of LAG3 expressing T cells in Ab1^fl/fl^ and SPC^ΔAb1^ mice. Results of unpaired two-tailed Student’s *t*-tests and Mann-Whitney tests comparing LAG3-expressing T cell populations between Ab1^fl/fl^ and SPC^ΔAb1^ mice. *P* values and associated *U* or *t* statistics with associated degrees of freedom (df) for each comparison are reported. Statistical results in this table are for the data depicted in Figure S7.

| Ab1<sup>fl/fl</sup> vs SPC<sup>ΔAb1</sup> Comparison Results |  |  |  |
| --- | --- | --- | --- |
| Experiment | Cell type | P-value | t, df |
| Homeostasis analysis | Lung CD4 %LAG3+ | 0.6918 | t = 0.4032, 17 |
|  | Lung CD8 %LAG3+ | 0.3662 | t = 0.9285, 17 |
|  | Spleen CD4 %LAG3+ | 0.4868 | t = 0.7109, 17 |
|  | Spleen CD8 %LAG3+ | 0.6578 | t = 0.4508, 17 |
| 9 days post IAV infection | Lung CD4 %LAG3+ | 0.8912 | t = 0.1403, 17 |
|  | Lung CD4 #LAG3+ | 0.9855 | t = 0.01860, 17 |
|  | Lung CD8 %LAG3+ | 0.4240 | t = 0.8336, 17 |
|  | Lung CD8 #LAG3+ | 0.7958 | t = 0.2658, 17 |

**Extended Data Table 3:** Mixed-effects model analysis results for weight loss after IAV infection. Results of repeated measures mixed-effects model (REML) analysis with Geisser-Greenhouse correction factoring in both genotype and day post IAV infection as variables, with post-hoc multiple comparisons with Sidak’s correction between the strains at each time point post infection. *P* values and associated *F* or *t* statistics with associated degrees of freedom (df) for the overall effects as well as for each post-hoc comparison are reported, calculated based on weights expressed as a percentage of initial weight. Data depicted in this table are in Figure 4a.

| <u>Overall model effects</u> |  |  |
| --- | --- | --- |
| **Factor** | ***P*-value** | ***F* (df.n, df.d)** |
| Time | <0.0001 | 177.4 (2.020, 188.5) |
| Genotype | 0.1696 | 1.916 (1, 94) |
| Genotype x Time | 0.0002 | 3.476 (10, 933) |

| <u>Post-hoc comparison results by day</u> |  |  |
| --- | --- | --- |
| **Day post-infection** | ***P*-value** | ***t*, df** |
| 5 | >0.9999 | 0.08273, 93.52 |
| 6 | >0.9999 | 0.4987, 91.57 |
| 7 | 0.9995 | 0.6777, 90.66 |
| 8 | >0.9999 | 0.06608, 81.36 |
| 9 | >0.9999 | 0.2905, 82.35 |
| 10 | 0.987 | 0.9875, 84.87 |
| 11 | 0.8544 | 1.414, 91.06 |
| 12 | 0.738 | 1.593, 92.94 |
| 13 | 0.3238 | 2.141, 91.81 |
| 14 | 0.6217 | 1.744, 92.33 |
| 15 | 0.5448 | 1.839, 92.95 |

**Extended Data Table 4:** Mixed-effects model analysis results for weight loss after SeV infection. Results of repeated measures mixed-effects model (REML) analysis with Geisser-Greenhouse correction factoring in both genotype and day post SeV infection as variables, with post-hoc multiple comparisons with Sidak’s correction between the strains at each time point post infection. *P* values and associated *F* or *t* statistics with associated degrees of freedom (df) for the overall effects as well as for each post-hoc comparison are reported, calculated based on weights expressed as a percentage of initial weight. Data depicted in this table are in Figure 4b.

| <u>Overall model effects</u> |  |  |
| --- | --- | --- |
| **Factor** | ***P*-value** | ***F* (df.n, df.d)** |
| Time | <0.0001 | 145.9 (2.652, 87.69) |
| Genotype | 0.0037 | 9.582 (1, 37) |
| Genotype x Time | <0.0001 | 3.707 (14, 463) |

| <u>Post-hoc comparison results by day</u> |  |  |
| --- | --- | --- |
| **Day post-infection** | ***P*-value** | ***t*, df** |
| 2 | 0.5271 | 2.051, 31.28 |
| 4 | 0.095 | 2.878, 36.78 |
| 5 | 0.1109 | 2.818, 35.97 |
| 6 | 0.5318 | 2.033, 36.94 |
| 7 | 0.4242 | 2.175, 36.79 |
| 8 | 0.741 | 1.767, 34.18 |
| 9 | 0.4465 | 2.153, 33.4 |
| 10 | 0.0429 | 3.215, 32.91 |
| 11 | 0.2627 | 2.447, 31.76 |
| 12 | 0.6094 | 1.95, 29.34 |
| 13 | 0.4114 | 2.221, 27.74 |
| 14 | 0.352 | 2.325, 24.9 |
| 15 | 0.2886 | 2.441, 23.79 |
| 16 | 0.4587 | 2.16, 26.21 |
| 17 | 0.5085 | 2.1, 24.41 |

**Extended Data Table 5:** Statistical test results for comparisons of PD-L1 expressing AT2s and PD1, CD44, and CD62L expressing T cells in IAV infected Ab1^fl/fl^ and SPC^ΔAb1^ mice. Results of unpaired two-tailed Student’s *t*-tests, Welch’s *t*-test, and Mann-Whitney test, comparing PD-L1-expressing AT2 cells, as well as bulk CD4^+^ T cells, CD44/CD62L CD4^+^ T cells, and PD1-expressing T cell populations between Ab1^fl/fl^ and SPC^ΔAb1^ mice 9 days post IAV infection. *P* values and associated *U* or *t* statistics with associated degrees of freedom (df) for each comparison are reported. Statistical results in this table are for the data depicted in Figure S9.

| Ab1<sup>fl/fl</sup> vs SPC<sup>ΔAb1</sup> Comparison Results |  |  |  |
| --- | --- | --- | --- |
| Experiment | Cell type | P-value | t or U, df |
| Lung T cell analysis | # CD4 | 0.2933 | t = 1.073, 26 |
|  | CD4 %CD44<sup>+</sup> CD62L<sup>-</sup> | 0.5433 | t = 0.6176, 26 |
|  | CD4 %CD62L<sup>+</sup> CD44<sup>-</sup> | 0.8360 | t = 0.2091, 26 |
|  | # CD4 PD1<sup>+</sup> | 0.3491 | t = 0.9535, 26 |
|  | CD4 %PD1<sup>+</sup> | 0.4146 | t = 0.8291, 26 |
|  | # CD8 PD1<sup>+</sup> | 0.8229 | t = 0.2261, 26 |
|  | CD8 %PD1<sup>+</sup> | 0.7667 | t = 0.2998, 26 |
| PD-L1 expression analysis | AT2s %PD-L1<sup>+</sup> | 0.6806 | U = 27 |

**Extended Data Table 6:**
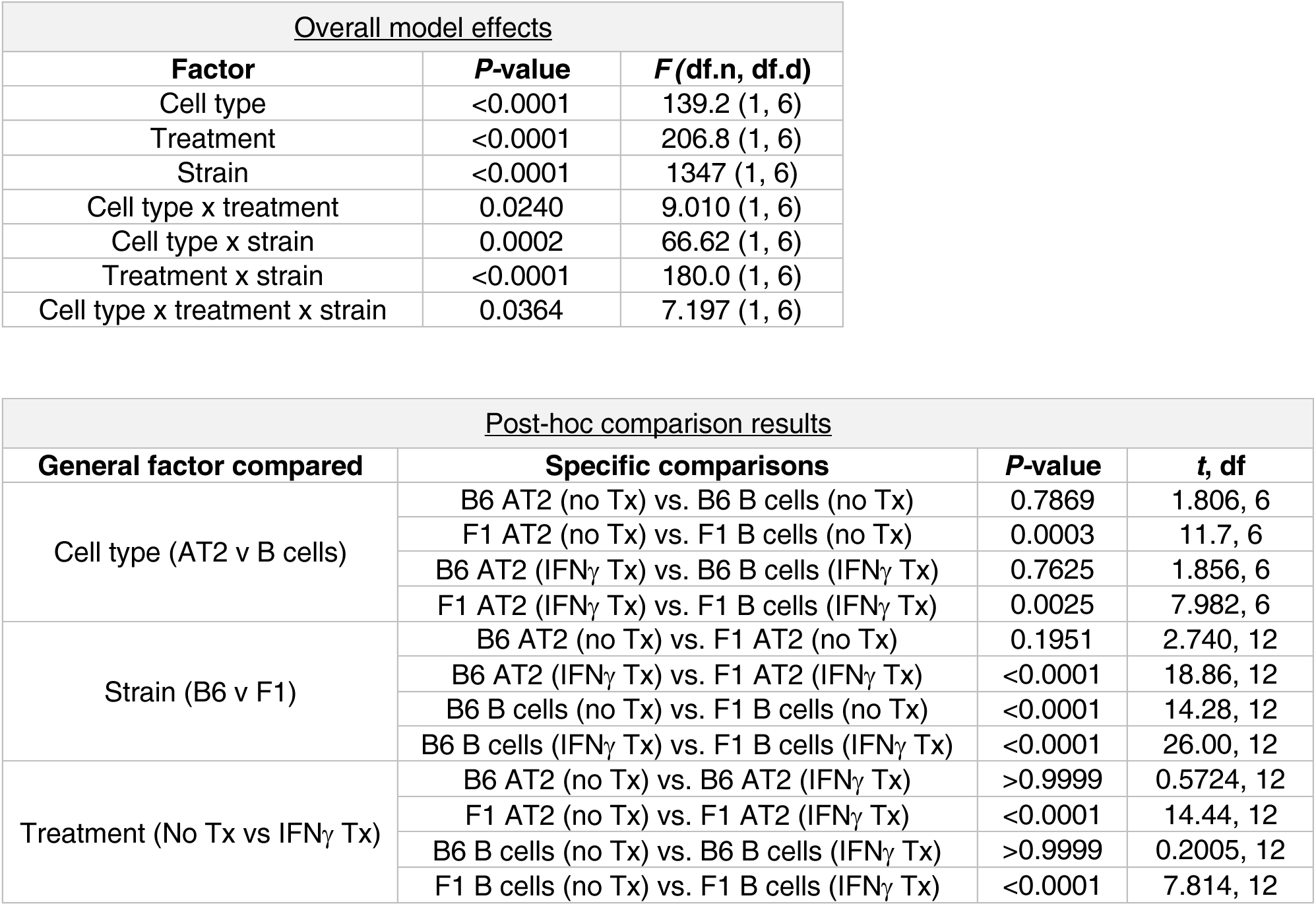
Three way ANOVA analysis results for YAe staining. Results of three-way ANOVA factoring in genotype, treatment (Tx), and cell type as variables, with post-hoc multiple comparisons with Sidak’s correction between means that differ by one factor only. *P* values and associated *F* or *t* statistics with associated degrees of freedom (df) for the overall effects as well as for each post-hoc comparison are reported. Data depicted in this table are in Figure 6c.

**Extended Data Table 7:** Two way ANOVA analysis results for I-A^b^ expression. Results of two-way ANOVA factoring in treatment and cell type as variables, with post-hoc multiple comparisons with Tukey’s correction between all means. *P* values and associated *F* or *q* statistics with associated degrees of freedom (df) for the overall effects as well as for each post-hoc comparison are reported. Data depicted in this table are in Figure 6d.

| <u>Overall model effects</u> |  |  |
| --- | --- | --- |
| **Factor** | ***P*-value** | ***F* (df.n, df.d)** |
| Cell type | <0.0001 | 357.8 (1,6) |
| Treatment | <0.0001 | 132.4 (1,6) |
| Interaction | <0.0001 | 146.5 (1,6) |

| <u>Post-hoc comparison results</u> |  |  |  |
| --- | --- | --- | --- |
| **General factor compared** | **Comparisons** | ***P*-value** | ***q*, df** |
| Cell type (AT2 v B cells) | F1 AT2 (no Tx) vs. F1 B cells (no Tx) | 0.0179 | 6.219, 6 |
| | F1 AT2 (IFNγ Tx) vs. F1 B cells (IFNγ Tx) | <0.0001 | 34.68, 6 |
| Treatment (No Tx vs IFNγ Tx) | F1 AT2 (no Tx) vs. F1 AT2 (IFNγ Tx) | <0.0001 | 23.61, 6 |
| | F1 B cells (no Tx) vs. F1 B cells (IFNγ Tx) | 0.9724 | 0.5985, 6 |
| Mixed | F1 AT2 (no Tx) vs. F1 B cells (IFNγ Tx) | 0.0077 | 7.411, 6 |
| | F1 AT2 (IFNγ Tx) vs. F1 B cells (no Tx) | <0.0001 | 30.42, 6 |

**Extended Data Table 8:** Two way ANOVA analysis results for I-E^d^ expression. Results of two-way ANOVA factoring in treatment and cell type as variables, with post-hoc multiple comparisons with Tukey’s correction between all means. *P* values and associated *F* or *q* statistics with associated degrees of freedom (df) for the overall effects as well as for each post-hoc comparison are reported. Data depicted in this table are in Figure 6e.

| <u>Overall model effects</u> |  |  |
| --- | --- | --- |
| **Factor** | ***P*-value** | ***F* (df.n, df.d)** |
| Cell type | <0.0001 | 4672 (1,6) |
| Treatment | <0.0001 | 1783 (1,6) |
| Interaction | <0.0001 | 1801 (1,6) |

| <u>Post-hoc comparison results</u> |  |  |  |
| --- | --- | --- | --- |
| **General factor compared** | **Comparisons** | ***P*-value** | ***q*, df** |
| Cell type (AT2 v B cells) | F1 AT2 (no Tx) vs. F1 B cells (no Tx) | <0.0001 | 23.66, 6 |
| | F1 AT2 (IFNγ Tx) vs. F1 B cells (IFNγ Tx) | <0.0001 | 123.9, 6 |
| Treatment (No Tx vs IFNγ Tx) | F1 AT2 (no Tx) vs. F1 AT2 (IFNγ Tx) | <0.0001 | 84.67, 6 |
| | F1 B cells (no Tx) vs. F1 B cells (IFNγ Tx) | 0.9986 | 0.2135, 6 |
| Mixed | F1 AT2 (no Tx) vs. F1 B cells (IFNγ Tx) | <0.0001 | 26.13, 6 |
| | F1 AT2 (IFNγ Tx) vs. F1 B cells (no Tx) | <0.0001 | 110.6, 6 |

**Extended Data Table 9:** Two way ANOVA analysis results for H2-Mαβ2 expression. Results of two-way ANOVA factoring in treatment and cell type as variables, with post-hoc multiple comparisons with Tukey’s correction between all means. *P* values and associated *F* or *q* statistics with associated degrees of freedom (df) for the overall effects as well as for each post-hoc comparison are reported. Data depicted in this table are in Figure 6g.

| <u>Overall model effects</u> |  |  |
| --- | --- | --- |
| **Factor** | ***P*-value** | ***F* (df.n, df.d)** |
| Cell type | <0.0001 | 8321 (1,6) |
| Treatment | 0.0006 | 43.37 (1,6) |
| Interaction | 0.0006 | 41.84 (1,6) |

| <u>Post-hoc comparison results</u> |  |  |  |
| --- | --- | --- | --- |
| **General factor compared** | **Comparisons** | ***P*-value** | ***q*, df** |
| Cell type (AT2 v B cells) | F1 AT2 (no Tx) vs. F1 B cells (no Tx) | <0.0001 | 89.17, 6 |
| | F1 AT2 (IFNγ Tx) vs. F1 B cells (IFNγ Tx) | <0.0001 | 94.75, 6 |
| Treatment (No Tx vs IFNγ Tx) | F1 AT2 (no Tx) vs. F1 AT2 (IFNγ Tx) | 0.0004 | 13.05, 6 |
| | F1 B cells (no Tx) vs. F1 B cells (IFNγ Tx) | 0.9998 | 0.1173, 6 |
| Mixed | F1 AT2 (no Tx) vs. F1 B cells (IFNγ Tx) | <0.0001 | 97.80, 6 |
| | F1 AT2 (IFNγ Tx) vs. F1 B cells (no Tx) | <0.0001 | 84.63, 6 |

